# A Genome-Wide Genetic Screen Identifies a Novel kDNA Replication Protein in Trypanosomes

**DOI:** 10.64898/2026.01.05.697628

**Authors:** Migla Miskinyte, Clirim Jetishi, Ana Kalichava, Alasdair Ivens, Martin Waterfall, Matt Gould, Lucy Glover, David Horn, Torsten Ochsenreiter, Achim Schnaufer

**Affiliations:** Institute of Immunology and Infection Research, University of Edinburgh, UK; Institute of Cell Biology, Faculty of Science, University of Bern, Switzerland; Wellcome Centre for Anti-Infectives Research, School of Life Sciences, University of Dundee, UK

## Abstract

Mitochondrial DNA of trypanosomatid parasites is organized into a topologically complex structure, named kinetoplast (kDNA). Replication, segregation and expression of kDNA involve an estimated ∼300 proteins, only a fraction of which have been identified and characterized. Here, we report the development of a genetic screen in *Trypanosoma brucei* to identify novel kDNA maintenance factors. Of the 20 highest-ranked genes identified, six are known kDNA maintenance factors. We selected one hit, *Tb927.8.4240*, a gene of previously unknown function, for experimental follow-up. Ultrastructure expansion microscopy using a tagged version of the protein reveals a dynamic localization during the cell cycle. RNAi-mediated ablation of *Tb927.8.4240* results in the progressive but incomplete loss of kDNA, with only a minor effect on the tripartite attachment complex, suggesting the protein is involved in kDNA replication but not segregation. The growth phenotype of *Tb927.8.4240* ablation is fully rescued in a kDNA-independent genetic background, confirming a specific role in kDNA replication.

In summary, we describe a functional genetic screen for the identification of kDNA maintenance factors in trypanosomes, validate one hit as a novel kDNA replication factor, and provide a prioritized hit list as a promising starting point for the future identification of additional factors.

## INTRODUCTION

Kinetoplastids are a group of flagellated protists that include the important human and animal pathogens of the genera *Trypanosoma* and *Leishmania* (1). A hallmark feature of these organisms is the kinetoplast DNA (kDNA), the organellar DNA within their single mitochondrion (2). Like mitochondrial DNA of other eukaryotes, kDNA encodes subunits of the mitochondrial oxidative phosphorylation (OXPHOS) system and the mitoribosome (3–5). However, unlike typical mitochondrial DNA, kDNA is arranged in a massive, catenated network comprising 5,000 – 10,000 minicircles and dozens of maxicircles, all physically linked into a planar disk (2, 6). The kDNA disk itself is physically linked to the flagellum’s basal body by the so-called tripartite attachment complex (TAC), a fibrous structure that crosses both mitochondrial membranes (7). The intricate kDNA network is not only visually striking when observed with electron or atomic force microscopy (8, 9) but is also essential for the survival and propagation of trypanosomes, as the encoded genes are critical for mitochondrial function and parasite viability (2, 5, 6, 10).

The complexity of the kDNA structure is functionally linked to equally complex mechanisms for mitochondrial gene expression. Out of the 18 mRNAs encoded in the maxicircles (20–30 kb in length in *T. brucei*), 12 require post-transcriptional editing by insertion and removal of uridines (11). For some mRNAs this editing is extensive and results in more than doubling of the mRNA length. The precise number of uridines to be inserted or removed, and the location of editing events within the mRNAs, is specified by short guide RNAs (gRNAs) by virtue of their complementarity to the fully edited sequence. The vast majority of gRNAs are encoded by minicircles (∼1 kb in length in *T. brucei*, with 3-4 gRNA genes per minicircle (12)). Hence, the sequence information for most kDNA-encoded genes is split between maxicircles and minicircles, and faithful replication and transfer to daughter cells of both types of molecules during cell division is therefore critical for parasite viability. Indeed, interference with kDNA maintenance is a key part of the mode of action of several anti-trypanosomatid drugs (13, 14).

Unsurprisingly, elucidation of the precise mechanism of kDNA replication and segregation, and of the molecular machineries responsible for these processes, has been a major focus of research, and the current state of knowledge of these processes has been detailed in several reviews (e.g. (2, 15, 16)). Replication and segregation of kDNA are highly regulated, multi-phase processes that occur once per cell cycle and involve the orchestrated action of numerous enzymatic and structural proteins and complexes. Summarised briefly, replication initiates with the release of individual minicircles from the kDNA network (presumably by topoisomerase 2) into the so-called kinetoflagellar zone (KFZ) between kDNA disk and inner mitochondrial membrane, where they undergo unidirectional theta-type replication (17). The process probably involves DNA polymerase IB and universal minicircle binding protein 1 (18, 19). The resulting daughter minicircles are then reattached to the network at so-called antipodal sites (APS) at opposite sides of the network periphery, a process that must require precise topological control to prevent entanglement and ensure network integrity (16). The APS are still poorly defined in composition and structure, but have been reported to contain several components of the kDNA replication machinery, including enzymes required for reattachment of minicircles into the network (topoisomerase 2 (20), possibly MiRF172 (21)), initiation of replication (primase 2 (22)), primer removal (helicases PIF1 and PIF5 (23, 24)), and gap repair (polymerase β (25), DNA ligase LigK-β (26)). The traditional model for minicircle replication postulates a sorting mechanism that ensures the distribution of the two daughter minicircles to opposite APS (2). More recently, reexamination of available localization data for kDNA replication factors led to an alternative ‘loose-diploid’ model that envisions two distinct lobes within the kDNA disk, each with an essentially complete set of minicircles that are released from the network laterally, replicated and then reattached to the same lobe (27). In both models, maxicircles remain attached to the network for replication (28), which likely involves DNA polymerases IC and ID (29, 30).

Following replication, the accurate segregation of the kDNA daughter networks into daughter cells is facilitated by the TAC, which links this process to the duplication and segregation of the basal body during cell division. The molecular architecture and biogenesis of the TAC has been dissected in considerable detail in recent years (reviewed in (15, 16)). TAC consists of three distinct subdomains: the extramitochondrial exclusion zone filaments (EZF) between the basal body and the outer mitochondrial membrane, the intramitochondrial unilateral filaments (ULF) between inner mitochondrial membrane and kDNA, and the differentiated membranes (DM) that bridge EZF and ULF. The DM subdomain is complex and consists of at least four different proteins (pATOM36, TAC40, TAC42 and TAC60; (15)). The EZF appears to consist of a single component, p197, that directly binds the basal body and is linked to the DM via TAC65 (31). The ULF consists of at least TAC53 (32), p166 (33) and TAC102 (34); the molecular details of its interaction with the kDNA disk itself remains to be elucidated but may involve at least two additional proteins that are not part of the TAC itself (35).

Overall, 63 different proteins have to date been reported as ‘kinetoplast-associated’ (36), which includes the kDNA replication factors and TAC components described above, and it was estimated that kDNA repair, replication and segregation may involve around 150 proteins (8). Important aspects of these processes that remain to be elucidated include the precise structural organisation of kDNA replication, including the mechanism of transport of minicircle replication intermediates to the reattachment sites, how topology and size of the network are controlled, and how replication of kDNA and of nuclear DNA are coordinated and integrated into the cell cycle.

The impressive progress that has been made in identifying components of kDNA replication and segregation in trypanosomes has resulted from a combination of bioinformatics, biochemical and imaging approaches, with each providing complementary insights into the molecular and cellular players and processes. In recent years, genome-wide RNAi screens in trypanosomes that select for phenotypes of interest after gene ablation have become a powerful tool in trypanosome research (37), but so far such screens have not been applied to the study of kDNA replication and segregation. Here, we developed an unbiased genetic screen for loss of kDNA after ablation of gene expression, aiming to identify novel factors that play critical roles in kDNA maintenance and its regulation and to help fill some of the gaps in understanding outlined above.

## MATERIALS AND METHODS

### *T. brucei* growth and whole-genome RNAi library generation

Bloodstream-form *T. brucei* 2T1^T7^ cells, resistant to blasticidin and puromycin, were cultured according to the published protocol (38). Cells were maintained in HMI-11 medium (39), supplemented with 10 fetal calf serum (FCS; Gibco) and 50 μg/ml penicillin/streptomycin (Thermo Fisher Scientific), under a 5% CO_2_ atmosphere, unless otherwise specified. A mutated F_1_F_O_-ATPase γ subunit (γL262P) was introduced into the 2T1^T7^ cell line by transfection with the NotI-digested pEnT6-NEO-γL262P plasmid, as described (40), and selected for resistance to neomycin at a concentration of 1 μg/ml. To confirm replacement of both alleles (γL262P/γL262P genotype), the relevant genomic region from selected transfectants was PCR-amplified and sequenced using primers 6 and 141 (**Table S3** lists all PCR and sequencing primers for this study). In contrast to the published procedure for generation of genome-wide RNAi libraries (38), expression of I-SceI meganuclease did not increase recombination and transfection efficiency in our cell lines. To obtain the desired library complexity, a total of 2.8×10^9^ 2T1^T7^-γL262P cells were transfected with 664 μg of the linearised pZJM RNAi library, using a total of 54 independent transfections (approximately 5.1×10^7^ cells and 12 μg of DNA per transfection). Following selection, transfected cells were grown in 300 ml of HMI-11 medium supplemented with 2 μg/ml phleomycin. After five days, cells were collected by centrifugation and resuspended in fresh HMI-11 medium with phleomycin, with selection continuing for an additional four days.

### Fluorescence activated cell sorting

Fluorescence activated cell sorting (FACS) was performed using a FACSAria cell sorter (BD Biosciences). *T. brucei* cells were grown under standard conditions at 37°C with 5% CO_2_, using HMI-11 medium with 10% FCS and antibiotics. To ensure cell vitality, the cultures were maintained in the exponential growth phase, not exceeding a density of 1×10^6^ cells/ml. During experimental procedures, cells were centrifuged at 1200 g for 5 minutes and then resuspended in Creek’s Minimal Media (CMM) (41) with an additional 10% FCS, followed by a recovery period of 2-3 hours at a lowered density of 5×10^5^ cells/ml. Post-recovery, the cells were labeled with dsFLUOR dye (Promega, E2671) for 2 minutes at a ratio of 1 μl dye per 4 ml of cell suspension. Preparing for fluorescence-activated cell sorting (FACS), the cells were treated with 10 μg/ml dihydroethidium (DHE), freshly made from a stock solution of 5 mg DHE in 500 μl DMSO, and incubated for 20 minutes at 37°C in the dark.

Subsequently, the cells were washed three times with CMM containing 4% FCS (1200g for 5 minutes), which had been pre-filtered through a 50 μm filter to eliminate any FCS aggregates that might affect the FACS process. Lastly, the cells were resuspended at a concentration of about 2×10^7^ cells/ml for sorting. To reduce nonspecific staining, sorting was conducted at a temperature of 10°C. Sorted cells were cultured for 80 hours to produce sufficient material for PCR amplification of RNAi inserts. Sorting was carried out for five independent biological replicates each, uninduced or induced for RNAi for 5 days.

### DNA deep sequencing

To amplify RNAi target fragments from the genomic DNA, we followed a previously described PCR protocol using LIB2f and LIB2r primers (38). After amplification, DNA was sent for Illumina deep sequencing (BGI, Hongkong; 100PE, HiSeq 4000). Paired end sequences were scanned for the presence of each primer subsequence (F=CCCCTCGAGG, R=ATCAAGCTTGGCC) and trimmed using a custom script to remove extraneous bases as appropriate. The resulting fasta output files for F, R and UNK (unknown) for each sample were aligned to the *T. brucei* TREU927 genome (Release 42 / v5.1 from TriTrypDB.org) using bowtie2 (version 2.2.7) (42) and parameters --very-sensitive -p 20 --no-unal. The resulting BAM files were sorted and indexed using SAMtools (version 1.12) (43), and alignment counts to predicted genes were generated from the BAM files with the appropriate bed file and BEDtools (v2.23.0) (44) using the multicov -bams *.bam command. The raw gene counts output from BEDtools was subsequently processed in R.

### RIT-seq data analysis

The statistical analysis of the RIT-seq data was performed in the R environment using edgeR package (45). The number of reads mapping to each CDS for each condition (5 days induced sorted, 5 days non-induced sorted and day 0 non-sorted populations; 5 replicates each) were normalized and log-transformed by the function calcNormFactors [trimmed mean of M-values (TMM) normalization].

### Construction of RNAi cell lines

The inducible stem-loop RNAi construct targeting gene Tb927.8.4240 was generated using the pQuadra system (46). Nucleotides 1-530 of the 8.4240 gene were amplified via PCR using primers 1722 and 1723 (**Table S3**). The oligonucleotides included specifically designed BstXI restriction sites to facilitate directional cloning (46). Ligation with BstXI-digested pQuadra1 and pQuadra3 plasmids generated pQuadra-8.4240, containing inverted repeats of the PCR product separated by a spacer region, which were confirmed by sequencing the plasmid using primers 116, 102 and 82 (**Table S3**). The NotI-linearized construct was then introduced via nucleofection (47) into *T. brucei* Lister 427 ’single marker’ bloodstream form cells, which express the T7 RNA polymerase and tetracycline repressor protein (48). Transgenic cell lines harboring the inducible RNAi cassette were selected with phleomycin. Subsequently, kDNA-independent (γL262P) and kDNA-dependent control (γWT) cell lines were constructed in this RNAi background by transfection with linearized pEnT6-PURO-γL262P and -γWT plasmids, respectively, as described previously (40).

### Quantitative real-time PCR

RNA from 1×10^8^ cells was isolated using the RNeasy Mini Kit (Qiagen). cDNA was synthesized from 1 μg of total RNA using the High-Capacity cDNA Reverse Transcription Kit (Applied Biosystems). Tb927.8.4240 RT-PCR was performed using 2 μl of a 1/10 dilution of cDNA, 10 μl of PowerUp SYBR Green Master Mix (Applied Biosystems), and 1 μl of 10 μM primer pair 1777/1778 (**Table S3**) using a Roche LightCycler. Analysis of the data was performed using the 2^-ΔΔCT method for relative quantification (49) with telomerase reverse transcriptase (TERT; Tb927.11.10190) as the reference gene (50) (primers 1358/1359, **Table S3**).

### Standard fluorescence microscopy

Standard fluorescence microscopy was performed as described previously (40). Briefly, cells were fixed with 2% (w/v) cold formaldehyde in phosphate buffered saline (PBS), washed once with PBS (1000 g, 5 min), and mounted on microscopy slides. DNA was visualized using Prolong Gold antifade reagent with 4′,6-diamidino-2-phenylindole (DAPI; Life Technologies), and images were captured using a Retiga 2000R Mono Cooled charged-coupled device camera attached to an Axioscope 2 or Axioimager Z2 (Carl Zeiss MicroImaging, Inc.) using either Plan-Apochromat 63x (1.40 NA) or Plan-Apochromat 100x (1.40 NA) phase-contrast objectives. The relative amount of DAPI-stained DNA was quantified using ImageJ software (51).

### Ultrastructure Expansion Microscopy (U-ExM)

Coverslips (12 mm, Epredia) were functionalized with poly-D-lysine (A3890401, Gibco) at room temperature for 30 min, followed by three washes with deionized water. Cells were then spread onto the coverslips and allowed to settle at room temperature for 10 min, fixed with 4% paraformaldehyde (PFA) for 10 min and permeabilized with 0.2% Triton X-100 in PBS for 5 min. Cells were washed twice with PBS and samples blocked with 4% bovine serum albumin (BSA) in PBS. Primary and secondary antibodies (see below for specifications) were diluted in PBS containing 2% BSA and were incubated for 45 min in between PBS washes. After the last wash with PBS, cells were anchored in PBS containing 0.7% formaldehyde (Sigma) and 1% acrylamide (AA; Sigma) at 37°C for 2 h. Gelation was performed using a freshly prepared solution composed of 19% sodium acrylate (Sigma), 10% AA, and 0.1% N,N’-methylenebisacrylamide (Sigma), supplemented with 0.5% ammonium persulfate (Thermo Fisher) and 0.5% tetramethylethylenediamine (Thermo Fisher). For gelation, 35 µl droplets of gelation solution were placed onto parafilm within a pre-cooled humidity chamber. Coverslips containing anchored cells were inverted onto droplets and incubated on ice for 5 min, followed by incubation at 37°C for 30 min to allow polymerization.

Gels were gently detached from the coverslips by incubating in 1 ml denaturation buffer (200 mM SDS, 200 mM NaCl, 50 mM Tris in deionized water, pH 9) with gentle shaking at room temperature for 15 min. Subsequently, gels underwent denaturation in the same buffer at 95°C for 30 min. After denaturation, gels were washed three times for 5 min each with PBS, then incubated for 1 h at room temperature with gentle agitation in PBS containing DAPI (5 µg/ml; Sigma) for DNA staining. Following staining, gels were expanded overnight in deionized water.

Antibodies were used as follows:

- anti-HA tag antibody, rabbit, 1:1000 (Thermo Fisher)
- Goat anti-Rabbit IgG (H+L) Cross-Adsorbed Secondary Antibody, Alexa Fluor™ 594, 1:1000 (Thermo Fisher)
- Mouse monoclonal anti-TAC102 (52), 1:1000
- Goat anti-Mouse IgG (H+L) Cross-Adsorbed Secondary Antibody, Alexa Fluor™ 488 (Thermo Fisher)
- Guinea pig anti-α-tubulin, 1:500 (ABCD Antibodies)
- Guinea pig anti-β-tubulin, 1:500 (ABCD Antibodies)
- anti-guinea pig IgG Alexa Fluor 647, 1:500 (Sigma-Aldrich)

Expanded gels were sectioned and mounted onto poly-D-lysine-coated 25 mm glass-bottom dishes (Cellvis, 35 mm dish with 20 mm microwell #1.5 cover glass). Z-stack imaging was performed using a Nikon Ti2 Kinetix Widefield Microscope equipped with a 100x objective (NA = 1.45). Imaging parameters included a z-step size of 0.2 µm and a pixel size of 65 nm. Images were deconvolved using Huygens HRM software and analyzed in ImageJ (51).

## RESULTS

### Development of a whole-genome scale RNAi screen for kDNA loss

We designed a screening approach based on a combination of genome-wide RNA interference, a novel kDNA staining protocol, and fluorescence-activated cell sorting (FACS) of kDNA^0^ cells; an overview is presented in **Fig. S1**. A key tool in our screening approach is the ability to genetically engineer bloodstream form *T. brucei* cells that are viable in the absence of kDNA (53). This is achieved by expressing a mutated copy of the γ subunit (γL262P) of the mitochondrial F_1_F_O_-ATPase, allowing for the creation of a mitochondrial membrane potential in the absence of kDNA-encoded genes (53). We transfected these kDNA-independent *T. brucei* cells with a whole-genome RNAi library (54, 55), achieving coverage of 60-87% for the eleven megabase chromosomes, based on Illumina sequencing of RNAi inserts PCR-amplified from transfected cells and mapping of reads to the *T. brucei* TREU927 reference genome (**Table S1**). Another essential advancement was our development of an *in vivo* DNA staining protocol that allowed the sorting, by flow cytometry, of trypanosome cells that had lost their kDNA. Initial attempts using dihydroethidium (DHE), a dye previously reported to be specific for kDNA (56), did not yield a sufficient separation of kDNA^+^ and kDNA^0^ cells by flow cytometry. However, subsequent trials of co-staining with DHE and dsFLUOR (57) resulted in a clear, if imperfect, separation of kDNA^+^ and kDNA^0^ *T. brucei* cells, as shown in **Fig. S1B**.

### Identification of putative kDNA maintenance factors

In previous studies, RNAi-mediated ablation of kDNA maintenance factors in bloodstream form parasites has been shown to result in kDNA loss after 3-5 days (35, 58). We therefore induced RNAi in our library cells for 5 days, after which kDNA^0^ cells were isolated by FACS (**Fig. S1A**). After extraction of total DNA from the kDNA^0^ cells, RNAi inserts were amplified by PCR (38) and sequenced on an Illumina platform (100 bp pair-ended) using an amplification-free library preparation protocol (sample ‘Day 5 +tet kDNA^0^’). For controls, we processed samples from (i) the full RNAi cell library prior to induction (‘Day 0’) and (ii) from cells un-induced for 5 days but sorted for kDNA^0^ (‘Day 5 -tet kDNA^0^’). This strategy aimed to distinguish gene fragments specifically enriched due to RNAi induction from those lost due to background noise or nonspecific effects. Illumina reads were mapped to a *T. brucei* reference genome (59) and differentially abundant reads per gene between tested conditions were determined.

**Fig. 1 panels A and B** show volcano plots, with thresholds for statistical significance set at log_2_ fold-change > 1.5 and *p* < 0.05. These criteria identified 1852 and 436 candidate genes, respectively, with an overlap of 132 genes between the two conditions (**Fig. 1C, detailed in Table S1**). While the precise number of genes required for kDNA maintenance is unknown, it is likely from the large number of hits identified under either condition that a considerable portion of the identified genes represent false positives. For example, only a small fraction of the genes from both hit lists encode proteins with experimentally determined mitochondrial localizations (36), contrary to what would be expected for most kDNA maintenance factors (**Table S2**). Specifically, the ‘Day 5 +tet kDNA^0^’ vs. ‘Day 0’ and ‘Day 5 +tet kDNA^0^’ vs. ‘Day 5 -tet kDNA^0^’ hit lists included only 249 (13.4%) and 85 (19.5%) genes encoding proteins with mitochondrial localization. Indeed, these percentages are similar to the percentage of genes encoding mitochondrial proteins (1,650, or 14.0%, according to the latest estimate (36)) among all 11,764 annotated genes in the reference genome (59). In addition, of the 63 genes reported to date (36) that encode kDNA-associated proteins, only 11 and 13, respectively, were included in our two hit lists (**Table S2**), suggesting false negatives as well. Notably, however, the 132 genes identified in both comparisons showed an enrichment for genes encoding mitochondrial proteins (32 genes, or 24%), including seven previously confirmed to be associated with the kinetoplast (36) (**Fig. 1C, Table S2**).

**Fig 1.**
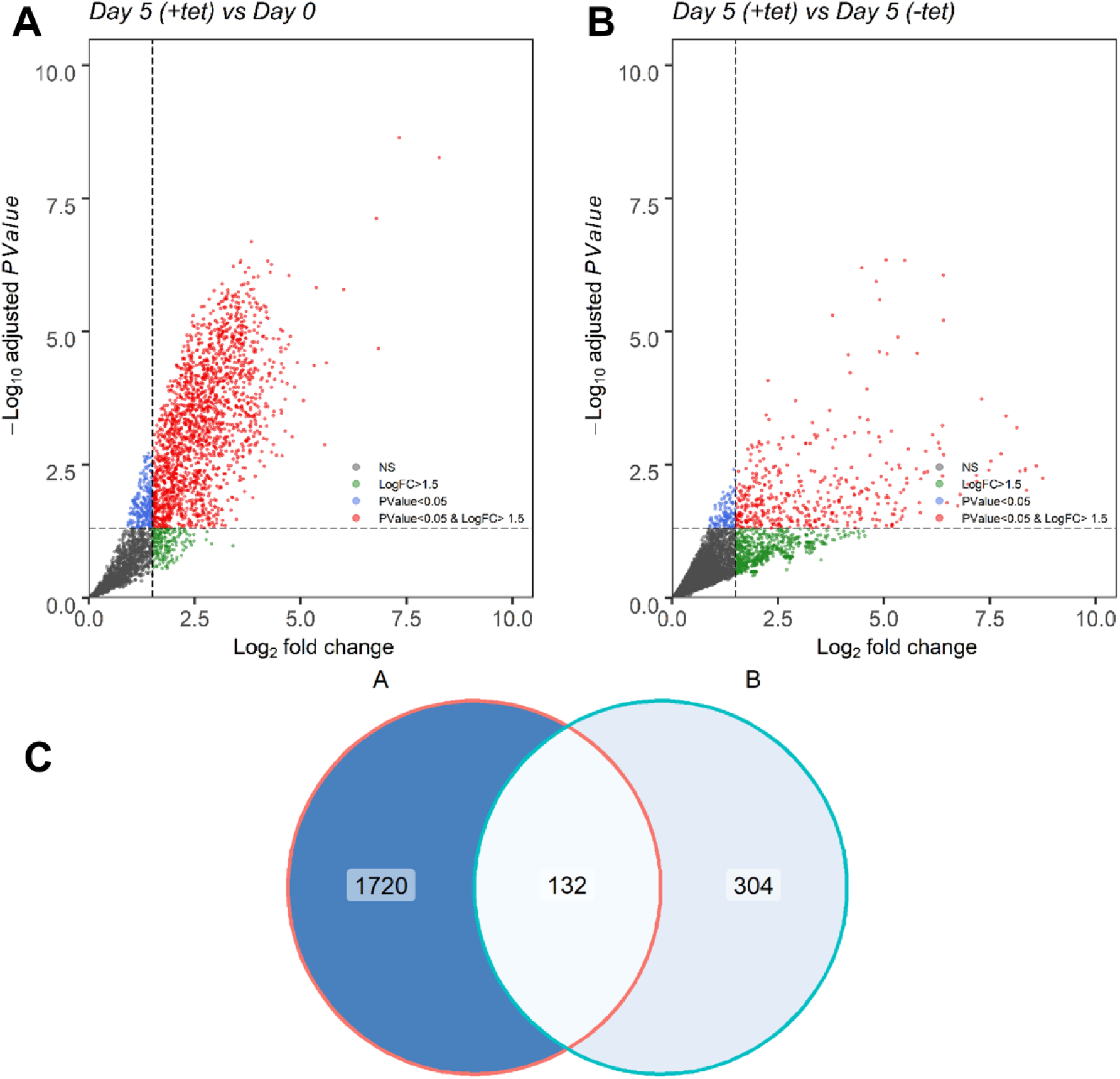
Identification of putative kDNA maintenance factors by genome wide RNAi screening. (**A, B**) Volcano plots showing the enrichment of genes in ’Day 5 +tet kDNA^0^’ compared to ’Day 0’ (A), and ’Day 5 -tet kDNA^0^’ (**B**). Significant enrichment is indicated for genes with a log_2_ fold change (log_2_FC) greater than 1.5 and an adjusted p-value of less than 0.05, shown in red. Each dot represents a gene; green dots represent non-significant (NS) genes with a log_2_FC less than 1.5 and/or an adjusted p-value equal to or greater than 0.05. The vertical dashed line represents the log_2_FC threshold of 1.5, and the horizontal dashed line indicates the p-value threshold of 0.05. Five biological replicates were used for all samples (n = 5). (**C**) Venn diagram representing the intersection of genes identified in both comparisons.

We therefore considered these 132 genes as our primary list of candidates. **Fig. 2** visualizes this selection, including published genome-wide data on localization (36, 60), the experimentally determined mitochondrial proteome of *T. brucei* (61–63), as well as currently known kDNA maintenance factors (16) (see also **Table S2**).

**Fig 2.**
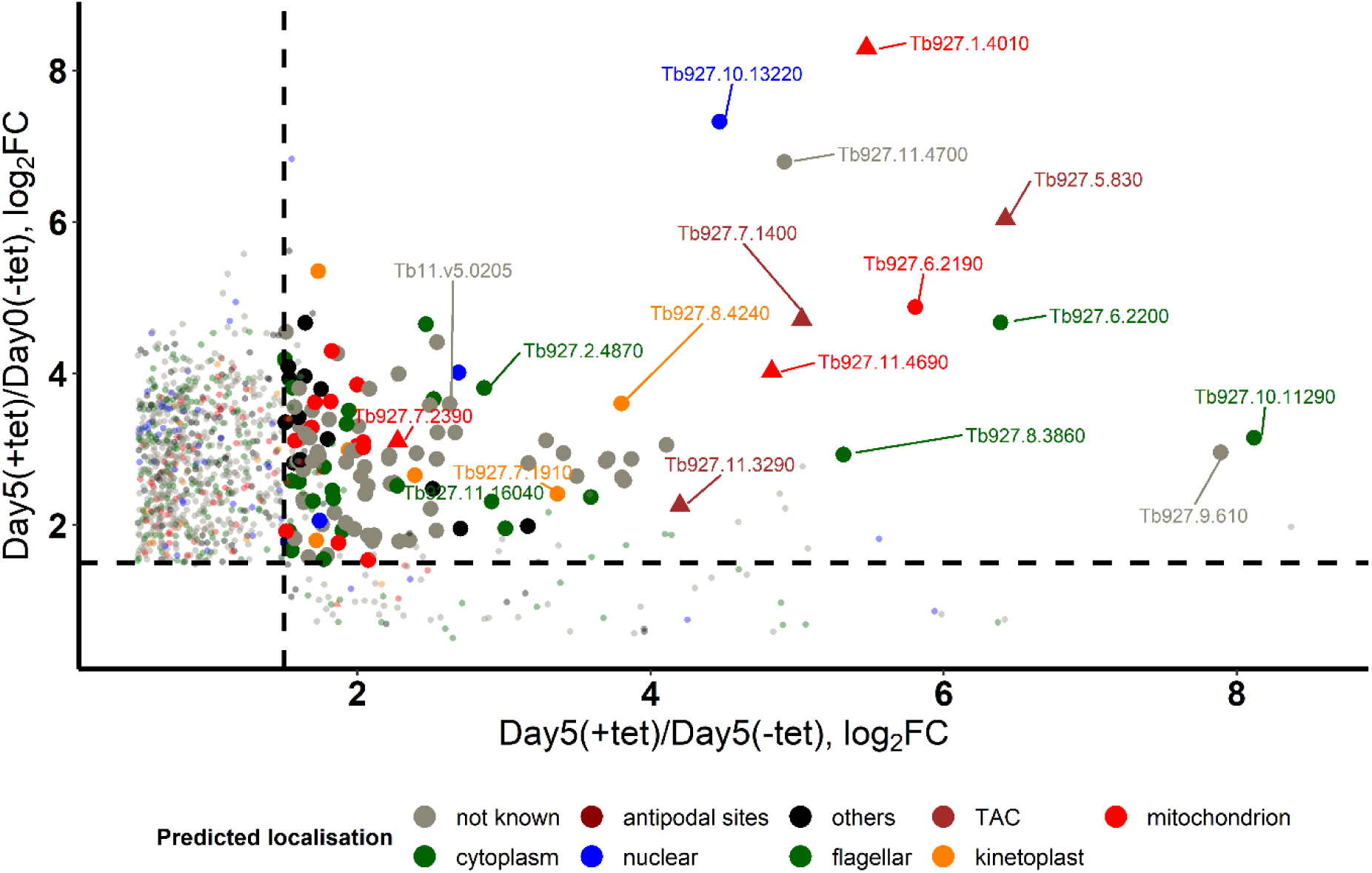
Visualization of primary set of 132 candidate genes implicated in kDNA maintenance, as identified through RNAi library screening. The visualized data include metadata on protein localization from TrypTag.org (36, 60), integration with published mitoproteome analyses (61–63), and previously identified kDNA maintenance factors (triangles; ref (16) and references therein). Statistically significant changes (adjusted *p*-value < 0.05) are indicated by larger circles. For clarity and reference, TriTrypDB gene identifiers are indicated for the top 20 ranked candidates based on the false discovery rate (FDR) in the y-axis comparison, detailed in Table 1 and Table S1.

**Table 1.** Top 20 candidates from the list of 132 primary candidates, ranked according to lowest false discovery rate (FDR) in the ‘Day 5 +tet kDNA^0^’ vs. ‘Day 5 -tet kDNA^0^’ comparison (Table S1).

| Rank | TriTrypDB ID | TriTrypDB Annotation | kDNA loss confirmed after knockdown | kDNA association confirmed | mitochondrial localization confirmed | Reference(s) (PMID) |
| --- | --- | --- | --- | --- | --- | --- |
| 1 | *Tb927.7.1400* | Tripartite Attachment Complex Protein 60 | yes | yes | yes | 29287109 |
| 2 | *Tb927.1.4010* | primase 2 | yes | yes | yes | 21257796 |
| 3 | *Tb927.10.13220* | cyclophilin type peptidyl-prolyl cis-trans isomerase, putative |  |  |  | 23556014 |
| 4 | *Tb927.5.830* | Tripartite attachment complex protein 65 | yes | yes | yes | 27436903 |
| 5 | *Tb927.11.4690* | mitochondrial DNA polymerase I protein B | yes | yes | yes | 12150917 |
| 6 | *Tb927.11.4700* | prostaglandin f synthase |  |  |  |  |
| 7 | *Tb927.8.4240* | hypothetical protein, conserved |  | yes | yes | 36804636, 32579605 |
| 8 | *Tb927.6.2200* | DJ-1 family protein, putative |  |  | yes | 37669165 |
| 9 | *Tb927.8.3860* | cytosolic iron-sulfur protein assembly 1 |  |  |  | 30346997 |
| 10 | *Tb927.6.2190* | hypothetical protein, conserved |  |  | yes | 37669165, 28485388 |
| 11 | *Tb927.11.3290* | Tripartite attachment complex protein 166 (p166) | yes | yes | yes | 18059470, 35709300 |
| 12 | *Tb927.9.16940* | variant surface glycoprotein (VSG, pseudogene), putative |  |  |  |  |
| 13 | *Tb927.11.16040* | prefoldin subunit, putative |  |  |  |  |
| 14 | *Tb11.v5.0467* | hypothetical protein, conserved |  |  |  |  |
| 15 | *Tb927.9.610* | hypothetical protein (pseudogene) |  |  |  |  |
| 16 | *Tb927.7.2390* | Tripartite attachment complex protein 102 | yes | yes | yes | 27168148 |
| 17 | *Tb927.10.11290* | hypothetical protein, conserved |  |  |  |  |
| 18 | *Tb927.7.1910* | pyridoxal phosphate containing glycine decarboxylase, putative |  |  | yes | 37669165 |
| 19 | *Tb927.2.4870* | kinetoplastid-specific dual specificity phosphatase, putative |  |  |  |  |
| 20 | *Tb927.6.1290* | hypothetical protein, conserved |  |  |  |  |

Encouragingly, when candidates were ranked based on the false discovery rate (FDR) for the ‘Day 5 +tet kDNA^0^’ vs. ‘Day 5 -tet kDNA^0^’ comparison, the top 20 included six genes encoding experimentally confirmed kDNA maintenance factors (four TAC components and two enzymes involved in replication): *TAC60* (64) (*Tb927.7.1400*, rank 1), *primase 2* (22) (*Tb927.1.4010*, rank 2), *TAC65* (65) (*Tb927.5.830*, rank 4), *mitochondrial DNA polymerase I protein B* (29) (*Tb927.11.4690*, rank 5), *p166* (33, 66) (*Tb927.11.3290*, rank 11) and *TAC102* (52) (*Tb927.7.2390*, rank 16) (**Table 1, Table S2**).

In addition to these six genes, four other candidates within the top 20 encode proteins with experimentally confirmed mitochondrial or kinetoplast locations (**Table 1**). *Tb927.8.4240* (rank 7) is annotated in TriTrypDB as a hypothetical gene, universally conserved among the sequenced kinetoplastids. A C-terminally tagged version of the 184 kDa protein has been localized to the kinetoplast (60). Notably, *Tb927.8.4240* is a paralog of the adjacent gene *Tb927.8.4230*, previously identified as encoding a putative interactor of LigK-β in a proximity labelling approach (67). LigK-β and LigK-α are kDNA-associated DNA ligases (26); only LigK-α expression has been successfully knocked down and shown to be essential for kDNA replication (26). While *Tb*927.8.4230 localizes exclusively at the antipodal sites, analogous to *T. brucei* LigK-β (26, 67), *Tb*927.8.4240 is reportedly localized in the proximity of kDNA, but not at the antipodal sites themselves (67).

*Tb927.6.2200* (rank 8) and *Tb927.6.2190* (rank 10) are neighboring genes, and close inspection of the mapped RNAi fragments revealed a fragment that included parts of both genes. It is therefore possible that only one of these two genes is a genuine hit, while the other is a ‘collateral’ hit. *Tb*927.6.2200 was predicted to be a mitochondrial protein based on its significantly reduced abundance in a mitochondrial fraction after ablation of ATOM40, a protein critical for mitochondrial protein import (63). The human homolog of this protein, DJ-1 (encoded by *Park7*), is also mitochondrial and is involved in protection from oxidative stress (68). *Tb*927.6.2190 is another ‘conserved hypothetical’ protein; its mitochondrial location was experimentally confirmed in multiple studies, and its expression appears to be regulated during the cell cycle (69). The final gene of interest in this subset, *Tb927.7.1910* (rank 18), encodes a component of the glycine cleavage complex (70), a crucial element in mitochondrial metabolism and function. This finding highlights the complex’s possible involvement in kDNA maintenance, potentially expanding the list of metabolic enzymes with such a dual function (71).

Six of the top 20 hits do not appear to encode mitochondrial proteins. *Tb927.10.13220* (rank 3) is annotated as a cyclophilin-type isomerase. Although some members of this family, like Cyclophilin D, have important mitochondrial functions (72), this gene has been predicted to encode a member of the Fam66 family of cell surface proteins (73). *Tb927.8.3860* (rank 9) has been experimentally confirmed as a component of the cytosolic iron-sulfur cluster assembly complex (74), which makes a putative association with kDNA maintenance unexpected. *Tb927.11.16040* (rank 13) is conserved among kinetoplastids, shows some homology to prefoldin-like chaperones and appears to be located in the cytoplasm. *Tb11.v5.0467* (rank 14), *Tb927.10.11290* (rank 17) and *Tb11.v5.0205* (rank 20), despite their conservation among kinetoplastids, possess uncertain cellular localizations (60), and lack identifiable functional motifs, which adds a layer of complexity to the functional characterization of these genes. Gene *Tb11.v5.0205* is identical in sequence to *Tb927.10.6130*. A tagged version of the encoded protein has been localized to the cytoplasm (60). The *Tb11.v5.0467* entry in TriTrypDB includes a 5’ UTR of nearly 3 kb that, in part, shares 100% identity with much of the *TAC102* coding sequence, suggesting that the contig that contains *Tb11.v5.0467* might be an assembly artefact and that this gene is a false positive hit.

Finally, the top 20 candidates include a further three genes that are probably false positive hits. *Tb927.916940* (rank 12) and *Tb927.9.610* (rank 15) are annotated as pseudogenes. Tb927.11.4700 (rank 6) is likely a ‘collateral hit’: it is located next to *Tb927.11.4690* (rank 5, see above) in the genome, and, similar to *Tb927.6.2200* and *Tb927.6.2190*, a close inspection of the mapped RNAi fragments revealed a fragment that included parts of both genes. Thus, the identification of *Tb927.11.4700* may result from its genomic proximity to a true positive hit rather than involvement in kDNA maintenance.

### Confirmation of *Tb*927.8.4240 as a novel protein involved in kDNA maintenance

Based on the above considerations, we prioritized *Tb927.8.4240* for follow-up studies. The 184 kDa protein product has no recognizable features or domains, except an intrinsically disordered region at its C-terminus (residues 1464-1689) predicted by InterPro (75). The structural model generated by AlphaFold (https://alphafold.ebi.ac.uk/entry/Q57W11) (76) has only a few domains of high confidence, with limited structural similarity to the following three proteins: an uncharacterized protein of *Etheostoma spectabile*, a fungal-specific transcription factor, and a LisH domain-containing *Anopheles* protein. As reported previously, *Tb927.8.4240* and its genomic neighbor in *T. brucei*, *Tb927.8.4230* (a gene encoding a smaller protein of 119 kDa) are the closest homologs of each other in the *T. brucei* genome (E-value 9e-41 in a protein BLAST) (67); they are likely paralogs that resulted from a gene duplication event. A search for potential orthologs and their genomics locations in trypanosomatids using the TriTryp database (www.tritrypdb.org) (77) confirms that both genes are highly conserved and usually syntenic in trypanosomatids. An exception is *T. cruzi*, where a gene specific for that species has been inserted between the two paralogs. Interestingly, in *T. congolense* IL3000, both genes – although still syntenic with their orthologs in other trypanosomatids, appear to have degraded and are annotated as pseudogenes *TcIL3000.A.H_000602900* and *TcIL3000.A.H_000603000* on chromosome 8.

Orthologs of *Tb927.8.4240* and *Tb927.8.4230* appear to be absent from the free-living genus *Bodo* (67), the closest known relative of trypanosomatids. A syntenic *Tb927.8.4240* ortholog is present in the early-branching trypanosomatid *Paratrypanosoma confusum*, but a *Tb927.8.4230* ortholog is missing in this species. This suggests a scenario where the gene duplication that gave rise to the two paralogs occurred after *P. confusum* had branched off; however, it is also possible that the *Tb927.8.4230* ortholog was secondarily lost in that species. **Fig. S2** shows sequence conservation for the orthologs from five selected trypanosomatids, *T. brucei*, *T. vivax*, *T. cruzi*, *L. major* and *Crithidia fasciculata*. Overall, within the *Trypanosoma* genus, *Tb*927.8.4240 and *Tb*927.8.4230 protein sequences are 57-59% and 43-47% identical, respectively (**Fig. S2A**). Areas of relatively high conservation are concentrated in six and two clusters, respectively (**Fig. S2B**; top and middle panel). Despite their common ancestry and homology, the calculated isoelectric points (pI) of the two paralogs are markedly different. While *Tb*927.8.4240 has a pI of 8.2, *Tb*927.8.4230 has a pI of 5.2 (note the abundance of negatively charged glutamic acid and aspartic acid residues, particularly in the C-terminal region of *Tb*927.8.4230; **Fig. S2B**, middle panel). In the alignment of the ten selected orthologs and paralogs from the five species, areas of relatively high conservation are restricted to four regions, the last three resulting from a fragmentation of the long, second region of conservation among the *Tb*927.8.4230 orthologs (Fig. S2B, bottom panel; Fig. S2C).

To confirm the role of *Tb*927.8.4240 in kDNA maintenance, we ablated its expression in bloodstream form *T. brucei* by inducible RNAi (**Fig. 3**). In two independent RNAi clones, after 3 days of induction, mRNA levels were reduced to ∼50% and ∼70%, respectively (**Fig. 3B**), which resulted in a marked reduction in the population’s growth rate approximately 5 days post-induction (**Fig. 3A**, grey and red dashed lines). Induction of RNAi in a kDNA-independent cell line (i.e. expressing γL262P) did not result in a growth phenotype (**Fig. 3A**, green lines), despite a comparable reduction in mRNA levels (**Fig. 3B**), confirming a kDNA-specific growth effect. After 5 days of RNAi induction, the percentage of cells in the population that had completely lost their kDNA had increased substantially: from ∼0.2% to >10% for 0K1N cells (i.e., cells with 1 nucleus but no kDNA) and from 0% to ∼1% for 0K2N cells (**Fig. 3C)**. In contrast, cells with replicated and segregated kDNA (2K1N and 2K2N) had decreased substantially. For cells that still possessed visible kDNA 5 days post-induction, a quantification of the kDNA-to-nucleus size ratio at the same time point revealed an overall reduction in average kDNA size and an increased variability in kDNA size (**Fig. 3D**). Even in the kDNA-independent RNAi cell line, which could be observed for a longer period after induction, the percentage of 0K1N cells remained relatively stable at 10% (**Fig. S2**). Depletion of *Tb*927.8.4240 in WT cells or in γL262P cells by RNAi had only a very minor effect on the amount or localization of the kDNA-proximal TAC component TAC102 (15, 52) (**Fig. 3E**), ruling out a role for *Tb*927.8.4240 in TAC assembly or structure.

**Fig 3.**
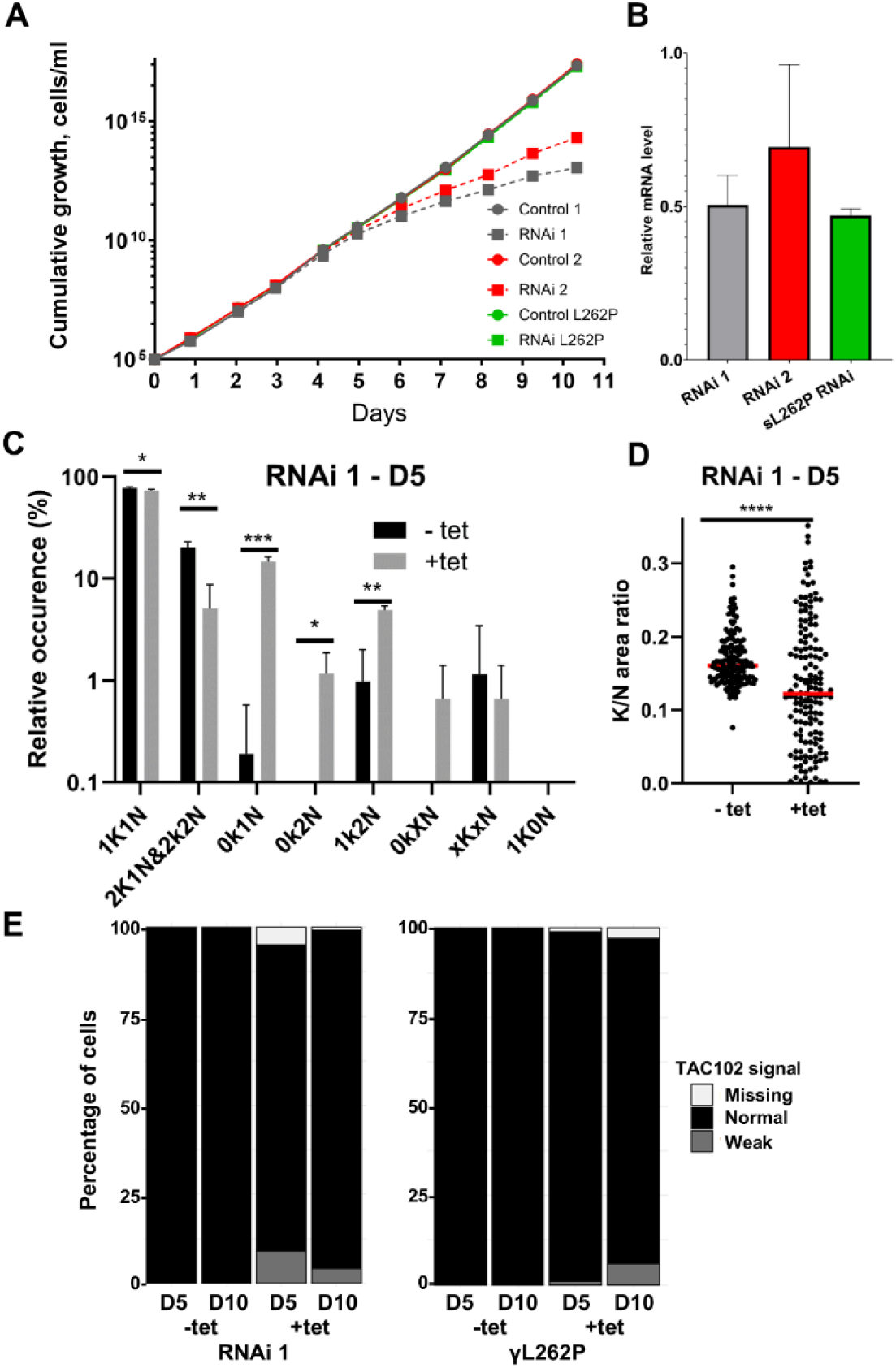
Phenotypic analysis of *Tb*927.8.4240 ablation in bloodstream form *T. brucei*. (**A**) Growth phenotype upon *Tb*927.8.4240 RNAi knockdown in WT cells (two independent RNAi clones (grey and red) is rescued in kDNA-independent cells (introduction of γL262P mutation; green). (**B**) Levels of mRNA measured by quantitative RT-PCR, at day 3 post induction (n = 3). (**C**) Quantification of relative occurrence of kDNA (K) and nuclei (N) in DAPI stained cells from induced (+tet) and non-induced (-tet) RNAi cell line 1 populations at day 5 post induction (n>100 cells for each triplicate; unpaired *t*-test P<0.05*, <0.01**, <0.001***). (**D**) The relative amount of kDNA to nucleus area in 1K1N RNAi cell line 1 at 5 days post-induction (Mann-Whitney test; P<0.0001****). (**E**) *Tb*927.8.4240 RNAi knockdown in WT cells or γL262P cells has only a minor effect on TAC102, as assessed with a TAC-specific antibody and epifluorescence microscopy (see Fig 4A; n>100 cells for each sample; paired *t*-test P<0.05).

Overall, these results suggested an important and specific role of *Tb*927.8.4240 in kDNA maintenance. The incomplete cessation of cell growth in kDNA-dependent cells and the incomplete kDNA loss in both kDNA-dependent and -independent cells could suggest some intrinsic variability in parasite dependence on *Tb*927.8.4240 function, or it could be a consequence of variable degrees of RNAi efficiency at the single cell level - the measured reduction in mRNA levels to 50-70% represents an average across the population.

### *Tb*927.8.4240 shows a dynamic localization at and near the kDNA APS and at the kDNA disk periphery

To confirm the association of the *Tb*927.8.4240 protein with kDNA, we expressed a hemagglutinin (HA) tagged version in procyclic form *T. brucei* and first determined the approximate location by fluorescence microscopy (**Fig. 4**). Co-staining for DNA and for TAC102 suggested that the *Tb*927.8.4240 protein is located at the APS.

**Fig 4.**
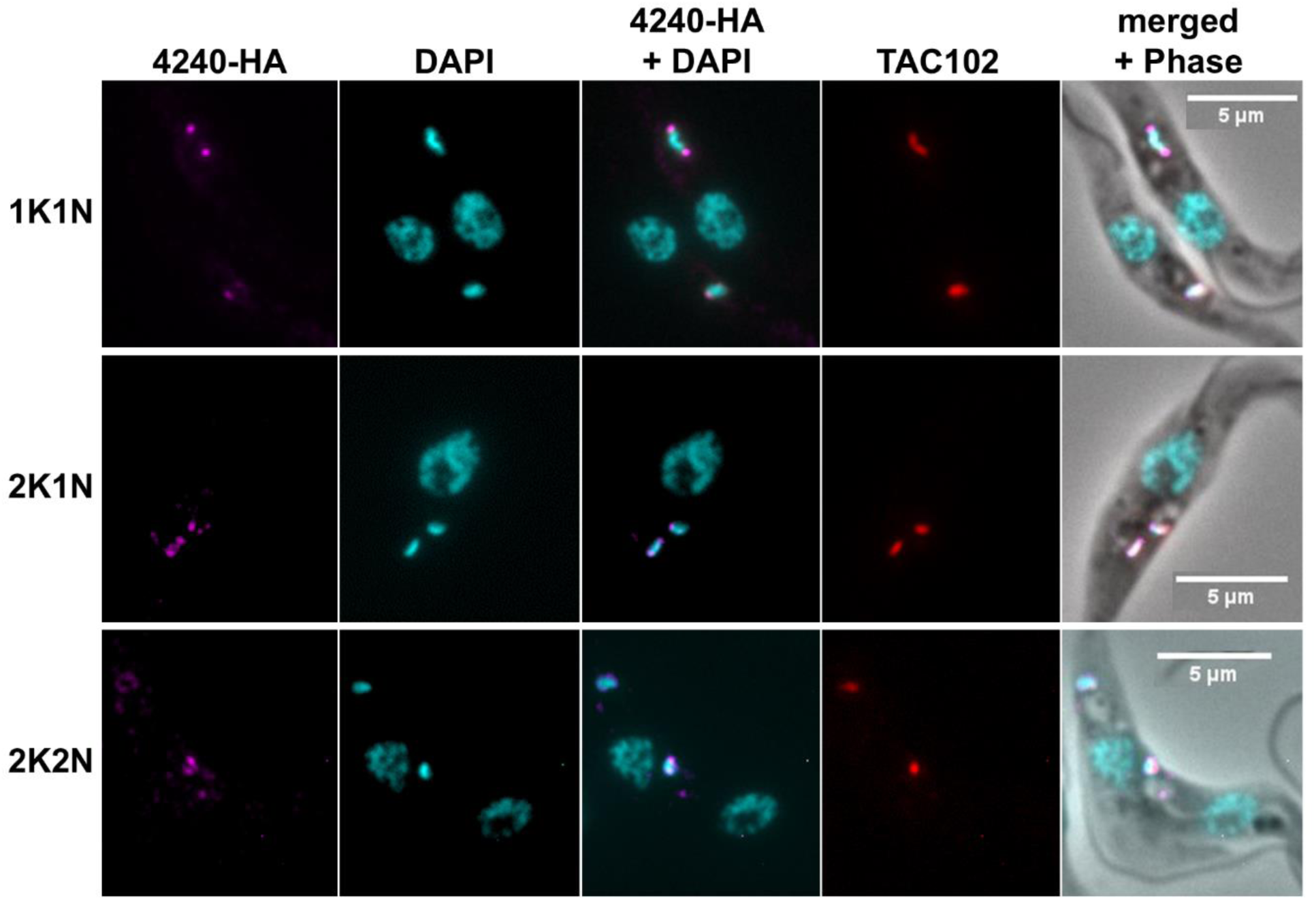
*Tb*927.8.4240 localization in procyclic *T. brucei* throughout the cell cycle assessed using standard epifluorescence microscopy. From left to right: *Tb*927.8.4240-HA tagged signal (magenta, anti-HA); nuclear and kDNA (cyan, DAPI); merge of anti-HA and DAPI; TAC102 (red, anti-TAC102); merge with phase contrast image. Scale bar 5 μm.

To precisely localize *Tb*927.8.4240, we performed Ultrastructure Expansion Microscopy (U-ExM; **Fig. 5**). Throughout the cell cycle, *Tb*927.8.4240 displayed distinct localization patterns when viewed from lateral and axial perspectives (**Fig. 5A**). In G1 / early S-phase cells (1K1N), we observed two punctate signals consistent with APS localization and clearly distinct from the TAC when viewed laterally (**Fig. 5A_i_**. **Fig. 5B**), while, when viewed axially, these signals either appeared at a punctate distribution around the edge of the disk (‘ring’) or in elongated shapes at opposite sites, presumably again reflecting APS localization (**Fig. 5A_ii,iii_**). These two distinct distributions were of similar frequency (**Fig. 5C**). In cells with dumbbell-shaped kDNA, i.e., in an advanced stage of replication, *Tb*927.8.4240 signals either exhibited similar elongated shapes with increased separation (**Fig. 5A_iv_**) or seemed shifted to oppose each other along the replicating kDNA rather than remaining antipodal (**Fig. 5A_v__-vii_**). The latter distribution was observed slightly more frequently (**Fig. 5C**). In segregating networks, the protein localization appeared as punctate rings again, with an additional localization along the ‘nabelschnur’ structure (78) between the disks (**Fig. 5A_viii_**).

**Fig 5.**
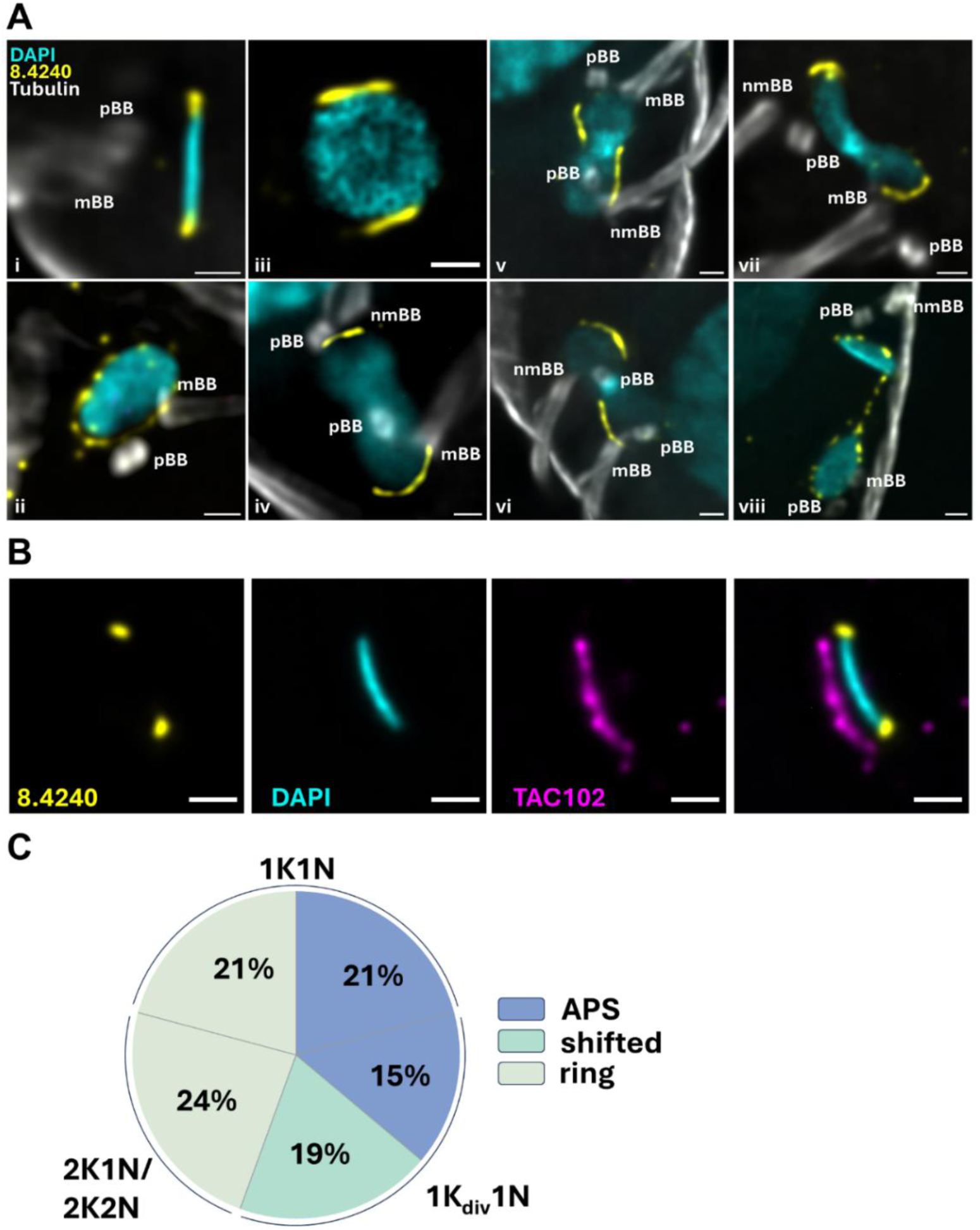
*Tb*927.8.4240 localization in procyclic *T. brucei* throughout the cell cycle, determined with Ultrastructure Expansion Microscopy (U-ExM). (**A**) *Tb*927.8.4240 localization pre replication (images i-iii), in replicating networks (images iv-vii), and in segregating networks (image viii), from both lateral (image i) and axial views (images ii-viii). *Tb*927.8.4240 is shown in yellow, kDNA in cyan, tubulin in grey; pBB, pro-basal body; mBB, mature basal body, nmBB, new mature basal body. Scale bar: 1 µm. (B) U-ExM image of *Tb*927.8.4240 (yellow) co-stained with TAC102 (magenta); kDNA is shown in cyan. Scale bar: 1 µm. (**C**) Quantitation of Tb927.8.4240 localization during the cell cycle as determined by U-ExM. Configuration 1K_div_1N represents cells with replicating kDNA, as in images iv-vii in panel A. A total of 144 cells were analyzed.

In summary, we found that the *Tb*927.8.4240 protein shows a dynamic, cell cycle-dependent localization.

## DISCUSSION

Biochemical and imaging approaches have resulted in substantial progress in the identification of genes involved in replication and segregation of the mitochondrial genome of trypanosomatids (16, 36, 79). Using a newly developed, genome-wide genetic screening approach, we have identified novel candidates for genes that are important in this process. We selected the gene with systematic ID *Tb927.8.4240*, annotated as a conserved hypothetical gene (www.TriTrypDB.org; (77)), for experimental follow-up.

Our bioinformatic analysis failed to identify any putative domains or motifs in the 184-kDa *Tb*927.8.4240 protein, except an intrinsically disordered region at its C-terminus. The structure predicted by AlphaFold is also not informative, as it has few domains of high confidence and very limited similarity to any domains with characterized function (76). Our gene expression knockdown by RNAi in bloodstream form *T. brucei* reduced the mRNA level to 50-70% and produced a slow growth phenotype after 5 days, a timescale that is consistent with growth phenotypes obtained in bloodstream forms for other kDNA maintenance factors (21, 80, 81). At day 5 post-induction, about 10% of cells had completely lost their kDNA (0K1N, or kDNA^0^), and a substantial reduction in kDNA size was observed for most remaining cells. Parasites then continued to grow at a slower rate, with no further increase in kDNA^0^ cells. It is uncertain why the percentage of kDNA^0^ cells did not eventually reach 100%, even in a kDNA-independent genetic background, but relatively mild growth and kDNA-loss phenotypes are not unusual after RNAi-mediated ablation in bloodstream form *T. brucei*, even for essential kDNA maintenance genes ((81) and references therein). The most straightforward explanations are the incomplete ablation of *Tb927.8.4240* mRNA (see above) and the known variability of RNAi efficiency within cell populations (82).

A tagged version of the *Tb*927.8.4240 protein was previously reported to localize near the kDNA antipodal sites (67). Our localization studies using tagged protein and standard fluorescence microscopy as well as U-ExM showed a dynamic, cell cycle-dependent location of the *Tb*927.8.4240 protein. For cells at the beginning of kDNA replication, we observed two alternative localizations within the population: either punctate, distributed around the rim of the kDNA disk, resulting in a ring-like appearance (Fig. 6, state *a*), or signals consistent with localization at antipodal sites (Fig. 6, state *b*). Viewed axially, APS localization was not dot-like, as in images obtained by conventional microscopy (60), but extended along the edge of the kDNA (**Fig. 6**, state *a*), similar to the recently discovered replication factor MiRF172 (21). In dumbbell-shaped (or bilobed) kDNA, characteristic of cells at an advanced stage of replication, the *Tb*927.8.4240 protein exhibited a similar localization pattern in ∼45% of cells (**Fig. 6**, state *c*), or its location was shifted along the edge of the elongated kDNA rather than remaining antipodal (**Fig. 6**, state *d*). It seems reasonable to assume that state *d* develops via state *c*, rather than directly from state *b*, as depicted in **Fig. 6**, although further experiments will be required to confirm this. This repositioning away from the antipodal sites is reminiscent of patterns observed after pulse labelling of replicating *T. brucei* minicircles with bromodeoxyuridine (BrdU), which led to the development of a model of *T. brucei* kDNA replication where back-and-forth oscillation of the kDNA disk facilitates a more equal distribution of daughter minicircles in the replicating network (83). Later metabolic labelling experiments with the thymidine analogue 5-ethynyl-2′-deoxyuridine (EdU) did not reproduce these patterns (84), and it was speculated that the DNA denaturation step required for BrdU antibody detection might have resulted in imaging artefacts, questioning the oscillation model (2).

**Fig 6.**
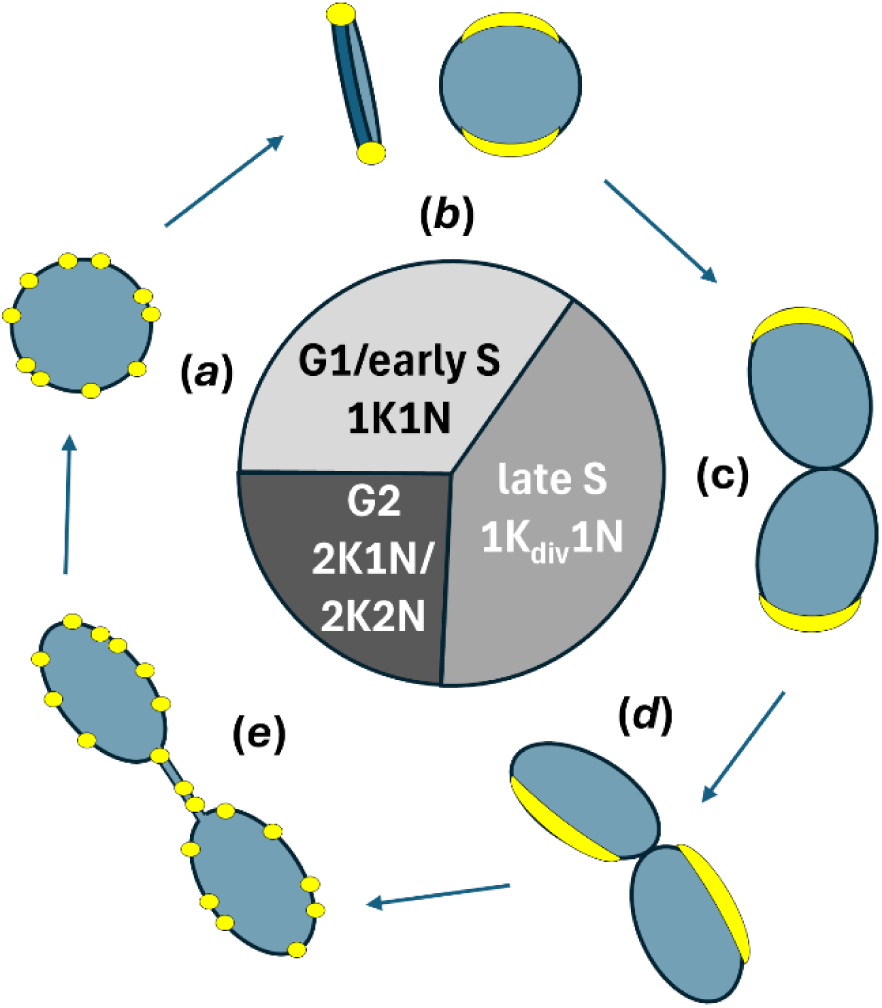
Model for dynamic localization of *Tb*927.8.4240 during kDNA replication. The model is based on the U-ExM studies shown in Fig. 5. Blue, kDNA; yellow, *Tb*927.8.4240 protein. Note that the respective size of the cell cycle stages in the diagram is drawn simply for convenience and not proportional to their actual duration. The three distinct states of localization are ring-like (a, e), antipodal (b, c) and shifted (d), in accordance with Fig. 5.

Interestingly, in segregating networks we also observed association of some *Tb*927.8.4240 protein with the ‘nabelschnur’ (German for umbilical cord) in addition to redistribution to the punctate pattern around the rim of the disk seen in state *a* (**Fig. 6**, state *e*). As the nabelschnur represents maxicircle DNA that is stretched out between the segregating networks (78, 85), this observation suggests that *Tb*927.8.4240 is not exclusively involved in minicircle replication. Previously, only two proteins have been reported that show association with this structure, the leucyl aminopeptidase *Tb*LAP1 (86) and *Tb*NAB70 (87). Similar to *Tb*927.8.4240, both proteins show co-localization with the kDNA disk during replication (specifically with the kDNA periphery in the case of *Tb*NAB70) and, additionally, with the nabelschnur during segregation. Phenotypes for ablation are clearly distinct from what we observed for *Tb*927.8.4240, though: ablation of TbLAP1 results primarily in accumulation of 2K2N cells (86), while ablation of TbNAB70 prevents kDNA segregation and results in accumulation of 1K2N cells (87).

There is additional precedence for changes in localization of kDNA replication factors during the cell cycle. For example, DNA polymerases IC and ID only accumulate at antipodal sites in actively replicating networks, but are dissipated throughout the mitochondrial matrix during other phases of the cell cycle (88, 89). Particularly relevant to our observation is the dynamic change in expression and localization of DNA LigK-α during the cell cycle in *C. fasciculata*, this enzyme being primarily observed on the faces of the kDNA disk in dividing cells, and having greater localization at the KFZ face in some cells and on the opposite face in others (90). *T. brucei* LigK-α is encoded by *Tb927.7.600*, the paralog of its neighboring gene, *LigK-β* (*Tb927.7.610*). The LigK-β protein interacts with the *Tb*927.8.4240 paralog *Tb*927.8.4230 (67), and both proteins are localized at the antipodal sites (26, 67). It is therefore tempting to speculate that *Tb*927.8.4240 dynamically interacts with DNA LigK-α, presumably specifically in dividing cells. The presence of a predicted intrinsically disordered region in Tb927.8.4240 is consistent with this idea, as the conformational malleability of these domains is well suited to facilitate dynamic interactions (91). This hypothesis needs to be tested in future experiments. Orthologs for both DNA ligases are also neighboring genes in the *B. saltans* genome (BSAL_42315 and BSAL_42320, www.TriTrypDB.org; (77)).

Bodonids are free-living kinetoplastids that are a sister group of the trypanosomatids, have free minicircles and lack a kDNA network structure (92, 93). It has been suggested that the network structure evolved in trypanosomatids to minimize minicircle loss during cell division (94). Our search of the TriTrypDB database for orthologs of *Tb927.8.4230* and *Tb927.8.4240* did not find orthologs for either gene in *B. saltans*. However, we identified a syntenic ortholog for *Tb927.8.4240*, but not *Tb927.8.4230*, in the early-branching trypanosomatid *P. confusum*. This suggests a scenario where recruitment of *Tb*927.8.4240 for kDNA replication and subsequent gene duplication and specialization of paralog function evolved with the need for more sophisticated kDNA replication and segregation during trypanosomatid evolution. Future studies should systematically investigate the presence and absence of known kDNA replication and segregation factors in the rapidly growing number of kinetoplastid genomes (95).

Above we described the validation and characterization of *Tb*927.8.4240 as a novel kDNA maintenance factor. This, along with the identification of six known kDNA maintenance factors in our top 20 list (**Table 1**) is proof-of-concept that, in principle, the genetic screen that we designed for identification of kDNA maintenance factors is effective. However, the following findings suggest that there is considerable room for improvement.

1. Of the most recent list of 63 known genes encoding kinetoplast-associated proteins, which include dozens of genes where ablation has been reported to result in disruption of kDNA biogenesis (16, 36), only seven are present in our list of 132 candidate genes (**Table S2**, **Fig. 1**). Thus, our screen produced a considerable number of false negatives.
2. The vast majority of proteins that are important for kDNA maintenance are expected to have mitochondrial localization. Although there is precedence for proteins that are involved in kDNA maintenance and that have an additional role outside of the mitochondrion (e.g. *mitochondrial topoisomerase 2*, *Tb927.9.5590* (53, 96)), and indeed we would expect some non-mitochondrial proteins with important roles in kDNA maintenance (e.g. for coordination of kDNA replication and segregation with the cell cycle), the number of proteins where this is the case is expected to be small. The 132 candidate genes include 32 genes (24%) where the protein product has been mapped to the mitochondrion (36). Although this is a clear enrichment over the estimated 14% of all genes that, according to a recent estimate (36), encode mitochondrial proteins, a percentage of 76% non-mitochondrial proteins suggests a considerable number of false positives in our list. Indeed, the list includes many genes with known functions that are not related to kDNA biogenesis. Thus, our screen clearly produced a considerable number of false positive hits.

How could the performance of the screen be improved to reduce the number of false negatives and false positives? One key step in our experimental design is separation of kDNA^+^ and kDNA^0^ cells by FACS (**Fig. S1**). Although this approach separated these cell types well, it was not perfect: in a control experiment with a pure population of kDNA^+^ cells, ∼0.5% of cells ended up in the kDNA^0^ gate (**Fig. S1B**). To produce enough genomic DNA for efficient amplification of RNAi inserts by PCR, we cultured cells for 80 hours after sorting (see Materials & Methods). Cells expressing γL262P tolerate kDNA loss well (53), but we noticed that in mixed populations of kDNA^+^ and kDNA^0^ cells, the percentage of the former slowly increases. Therefore, it would be beneficial to further optimize the separation of kDNA^+^ and kDNA^0^ cells by FACS and, also, the PCR protocol to allow efficient amplification of RNAi inserts directly after sorting. Finally, increasing the number of biological and technical replicates (we had used five biological replicates for the entire procedure from RNAi induction over sorting to library preparation and sequencing) would improve the identification of false hits with statistical methods.

Nonetheless, in addition to the characterized hit *Tb*927.8.4240, our hit list includes promising candidates for prioritization in future studies. This includes three candidates in our top 20 list with confirmed mitochondrial localization of the encoded proteins (*Tb927.6.2200*, *Tb927.6.2190*, *Tb927.7.1910*; **Table 1**) and 22 candidates further down in the list that also encode mitochondrial proteins (**Table S2**). Of particular note is *Tb927.5.520*, a gene encoding a putative stomatin-like protein, as proteins with the stomatin/prohibitin/flotillin/HflK/C (SPFH) domain have been reported as parts of complexes involved in maintenance of mitochondrial DNA (97).

In conclusion, we present an experimental strategy for a genetic screen that (i) has identified a novel kDNA maintenance factor, *Tb*927.8.4240, (ii) suggests further promising candidates for genes with important roles in kDNA maintenance, and (iii) points the way for second-generation screens with improved performance. Our characterization of *Tb*927.8.4240 suggests that the oscillation model for kDNA replication in *T. brucei* should be revisited, and that detailed and systematic phylogenetic studies of the evolution of known kDNA maintenance factors in kinetoplastids will be a feasible and fruitful endeavor.

## DATA AVAILABILITY

Raw sequencing reads have been deposited at the NIH SRA under BioProject ID PRJNA1402727.

## SUPPLEMENTAL DATA

SUPPLEMENTAL figures S1-S3 Tables S1-S3

## ACKNOWLEDGMENTS

This work was supported by Senior Non-Clinical Fellowship MR/L019701/1 from the UK Medical Research Council to A.S. and an Institutional Strategic Support Fund (ISSF3) award (reference IS3-R2.28) to A.S. for a salary for M.M. and consumables. Furthermore, the work was supported by a Swiss National Science Foundation grant (number 207525) and a grant from the Uniscientia foundation to T.O. and by a Wellcome Investigator Award (217105/Z/19/Z) to D.H..

## Supplemental Figures

**Fig S1.**
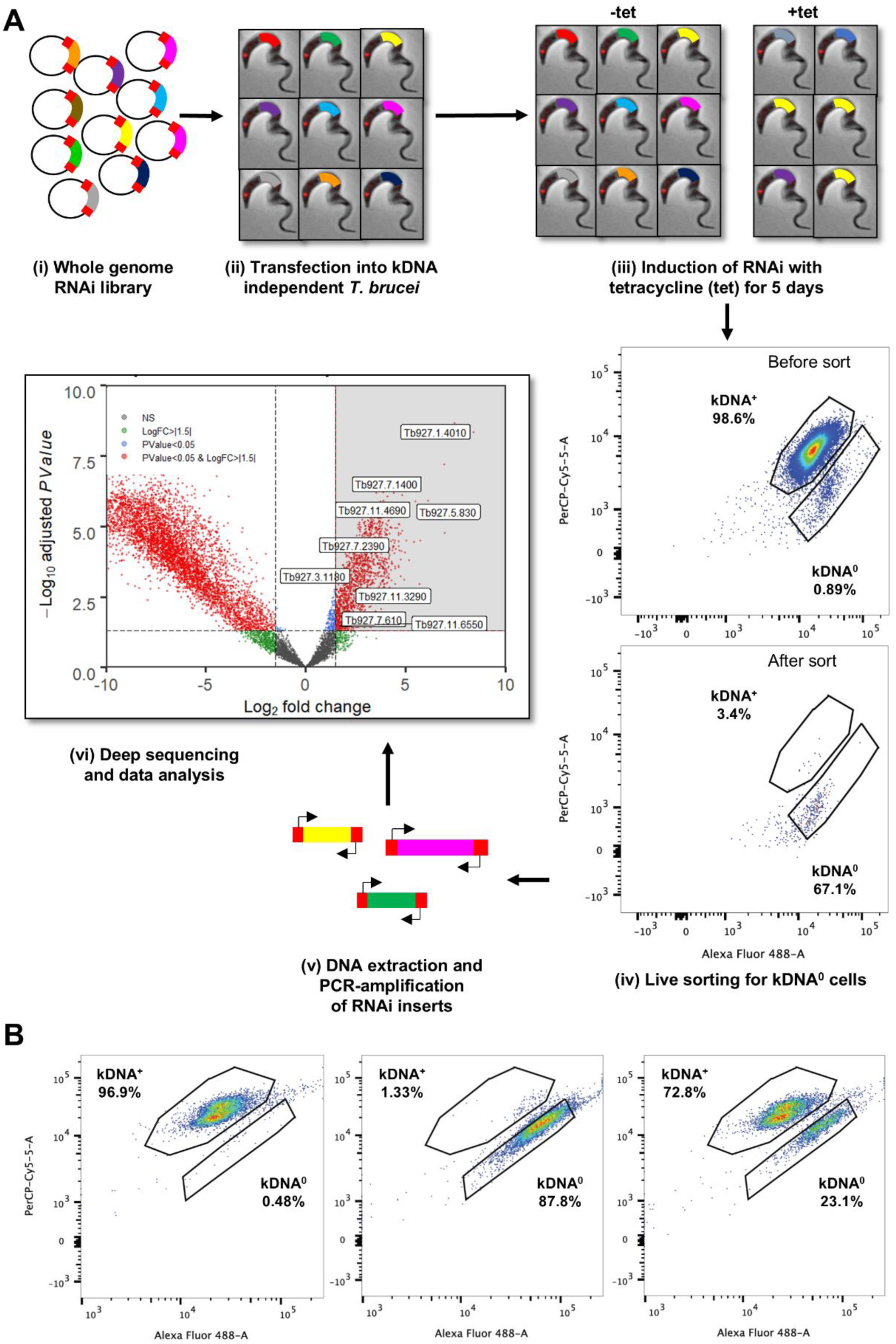
Screening approach. (**A**) Overview of the kDNA RIT-seq experimental design to find genes that are important for kDNA maintenance. (i, ii) A whole genome RNAi library (Alsford et al., Genome Research 2011) was transfected into kDNA-independent *T. brucei*. (iii) Upon RNAi library induction (+tet), fragments corresponding to genes essential for kDNA maintenance are expected to result in the loss of kDNA from the cell. Note that induction of some RNAi fragments will result in a growth defect and depletion of the cells from the population, but in the kDNA-independent background this should not be the case for genes of interest. (iv) To identify these genes, 5 days after induction, live cells are sorted by FACS after staining of DNA with two dyes, dihydro-ethidium (DHE) and dsFLUOR. These dyes differ in their affinities for kDNA vs. nuclear DNA, and measuring fluorescence in the Alexa Fluor 488 and Cy5-5 channels separates kDNA^+^ and kDNA^0^ populations. The representative image shows the population before sorting for one of the induced replicates. Isolated kDNA^0^ cells are then scaled up by growing them for 80 hours in culture media before extraction of total DNA. (v) RNAi fragments are PCR-amplified using RNAi cassette–specific primers. (vi) PCR amplicons are then fragmented and Illumina sequenced. After mapping to the reference genome, the number of mapped reads per gene is compared for the following samples: Comparison 1: day 5 +tet, sorted for kDNA^0^ (n = 5) vs. uninduced day 0 (n = 5), unsorted. Comparison 2: day 5 +tet, sorted for kDNA^0^ vs. day 5 -tet, sorted for kDNA^0^ (n = 5). (**B**) Establishing the FACS protocol for differential staining of kDNA^+^ and kDNA^0^ cells. Images show the FACS plots from optimized conditions using 1 μl dsFLUOR dye per 4 ml of cell suspension and 10 μg/ml dihydroethidium (DHE; see Materials and Methods for details). From left to right, cell populations are: control (kDNA^+^) populations of kDNA-independent *T. brucei*, the same cell line after treatment with ethidium bromide to remove kDNA (kDNA^0^), and a 1:1 mix of these populations.

**Fig S2.**
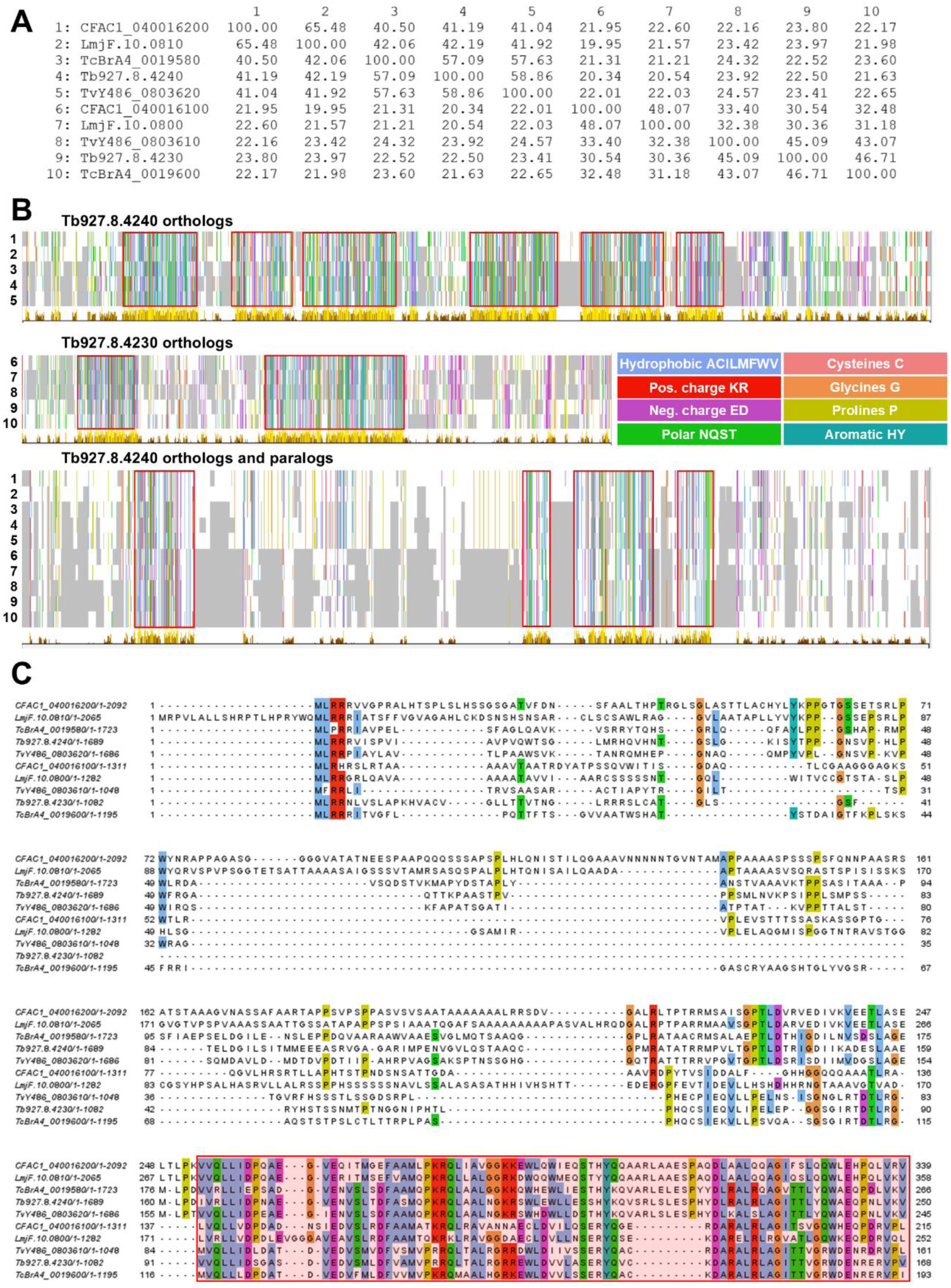
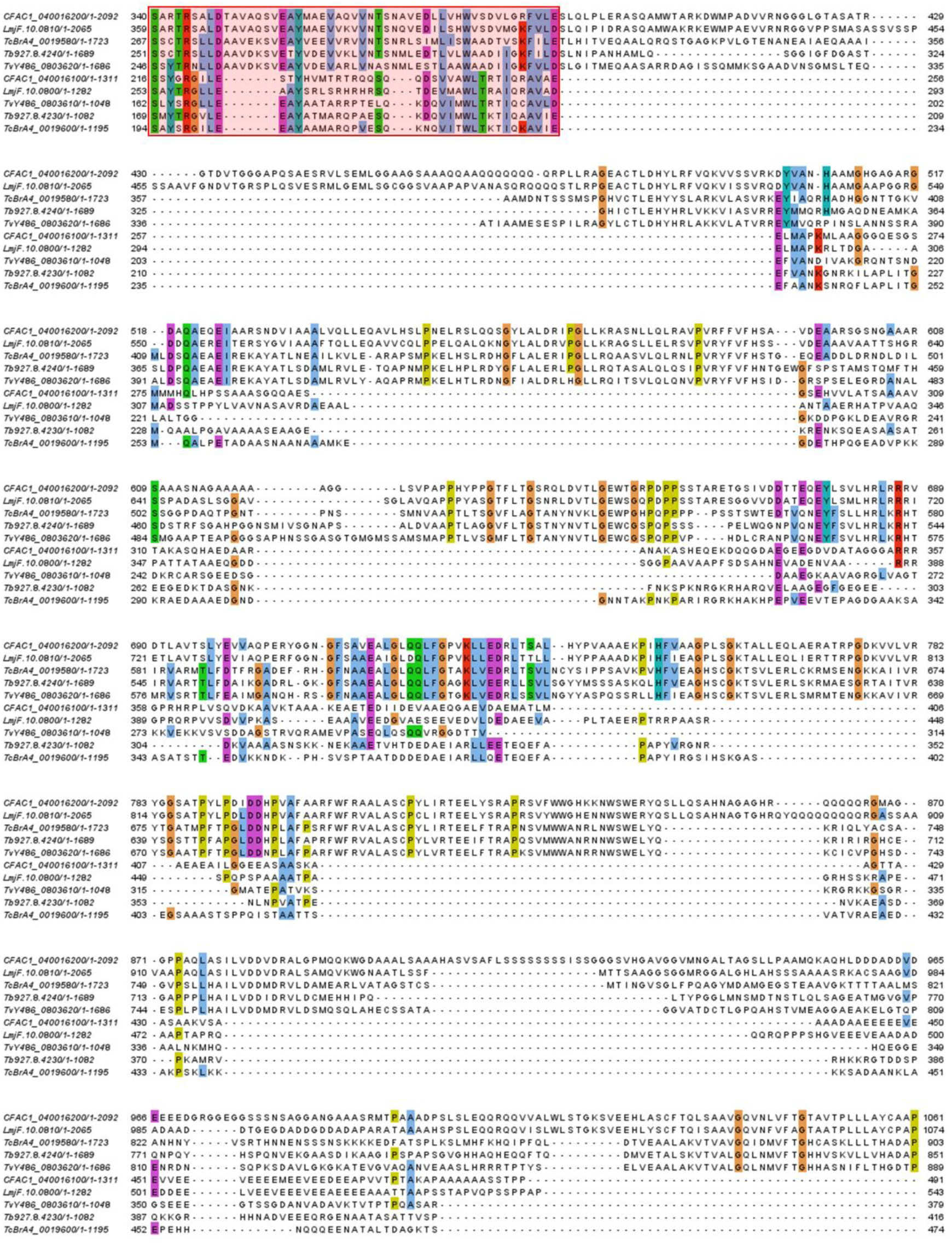
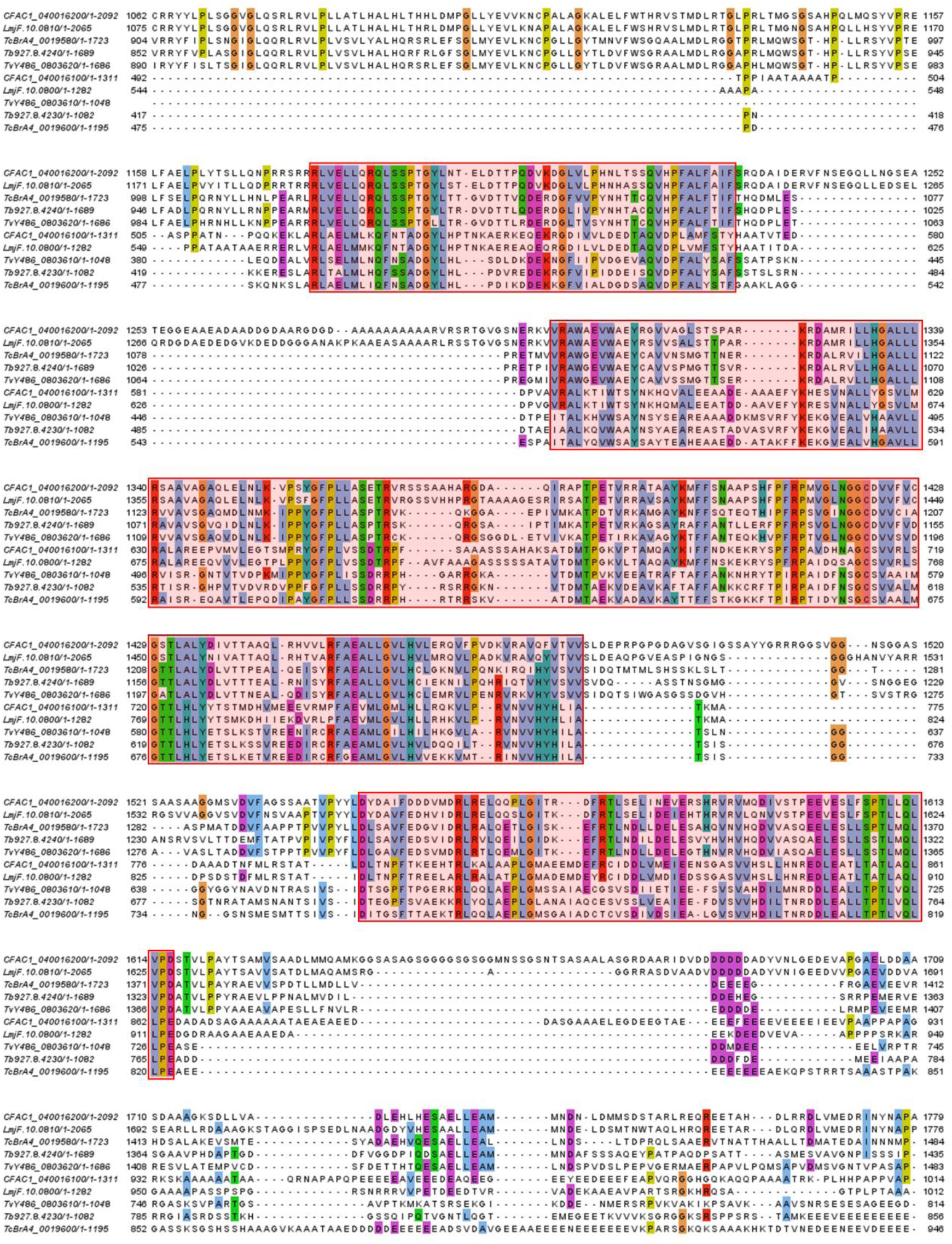
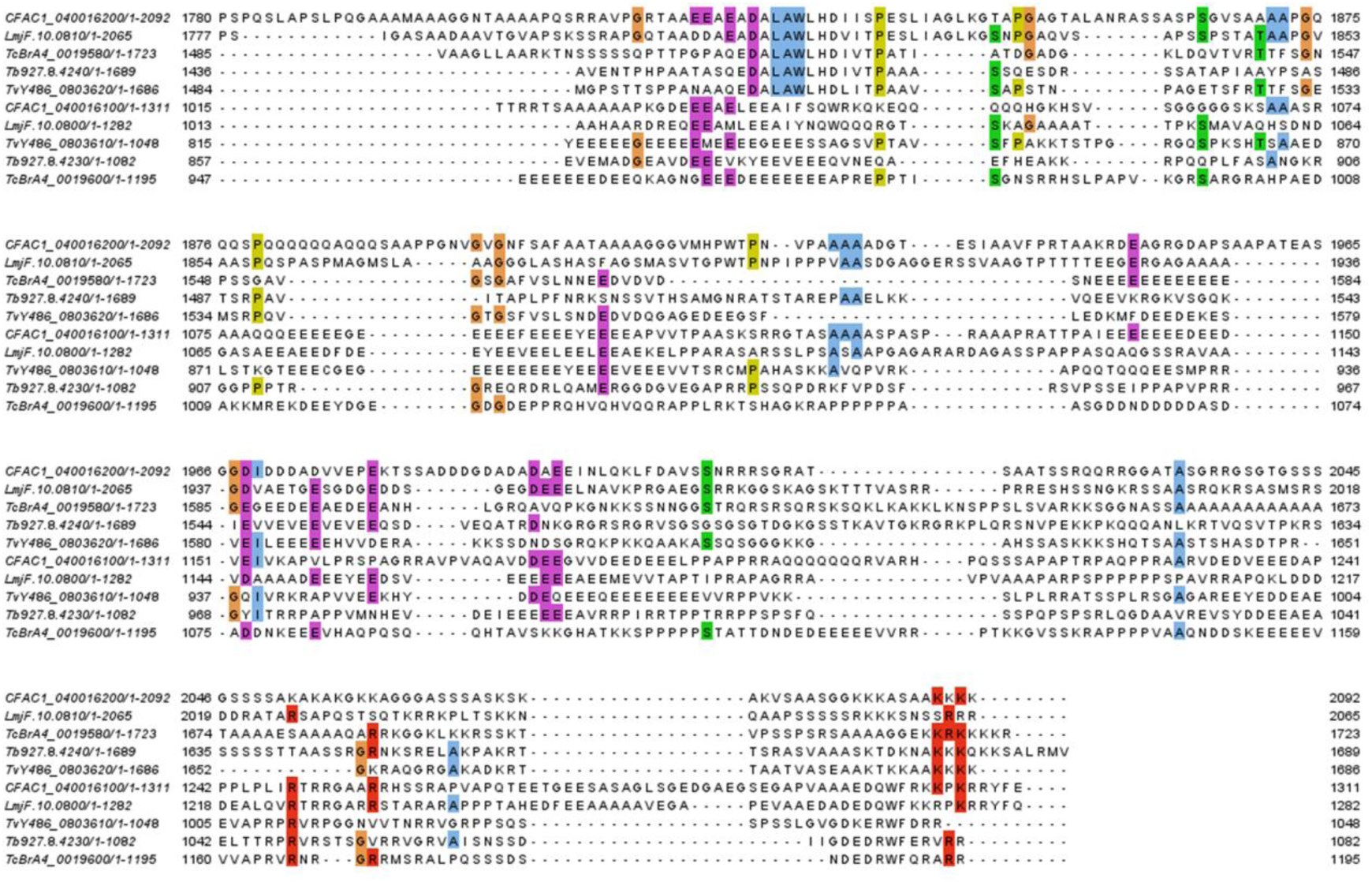
Sequence conservation of *Tb*927.8.4240 protein orthologs and paralogs in selected trypanosomatids. Sequences 1-5: *Tb*927.8.4240 orthologs. Sequences 6-10: Tb927.8.4230 orthologs. 1, 6: *Crithidia fasciculata* (Cf-C1). 2, 7: *L. major* (Friedlin). 3, 8: *T. cruzi* (Brazil A4). 4, 9: *T. b. brucei* (TREU 927). 5, 10: *T. vivax* (Y486). (**A**) Percent identity matrix, generated using Clustal 2.1. (**B**) Overviews of sequence conservation in *Tb*927.8.4240 orthologs (top panel), *Tb*927.8.4230 orthologs (middle panel) and *Tb*927.8.4240 orthologs and paralogs combined. Alignments were generated using Muscle (https://www.ebi.ac.uk/jdispatcher/msa/muscle) (98) and visualized using Jalview (99). Amino acids are colored according to the Clustal color scheme in Jalview. Each residue in the alignment is assigned a color if the amino acid profile of the alignment at that position meets some minimum criteria specific for the residue type (40% identity threshold). Blue: hydrophobic (A, C, I, L, M, F, W, V). Red: positive charge (K, R). Magenta: negative charge (E, D). Green: polar (N, Q, S, T). Pink: cysteines (C). Orange: glycines (G). Yellow: prolines (P). Cyan: aromatic (H, Y). White: unconserved. A track with bars underneath the individual sequences reflects the conservation of the physico-chemical properties for each column of the alignment, calculated based on the AMAS method of multiple sequence alignment analysis (100). The higher the score, the taller the bar and the lighter the shading (highest scores being reflected by yellow). (**C**) Detailed sequence alignment of selected Tb927.8.4240 orthologs and paralogs. Sequences and color codes are as in panels A and B. Red boxes and shading indicate regions of relatively high conservation.

**Fig S3.**
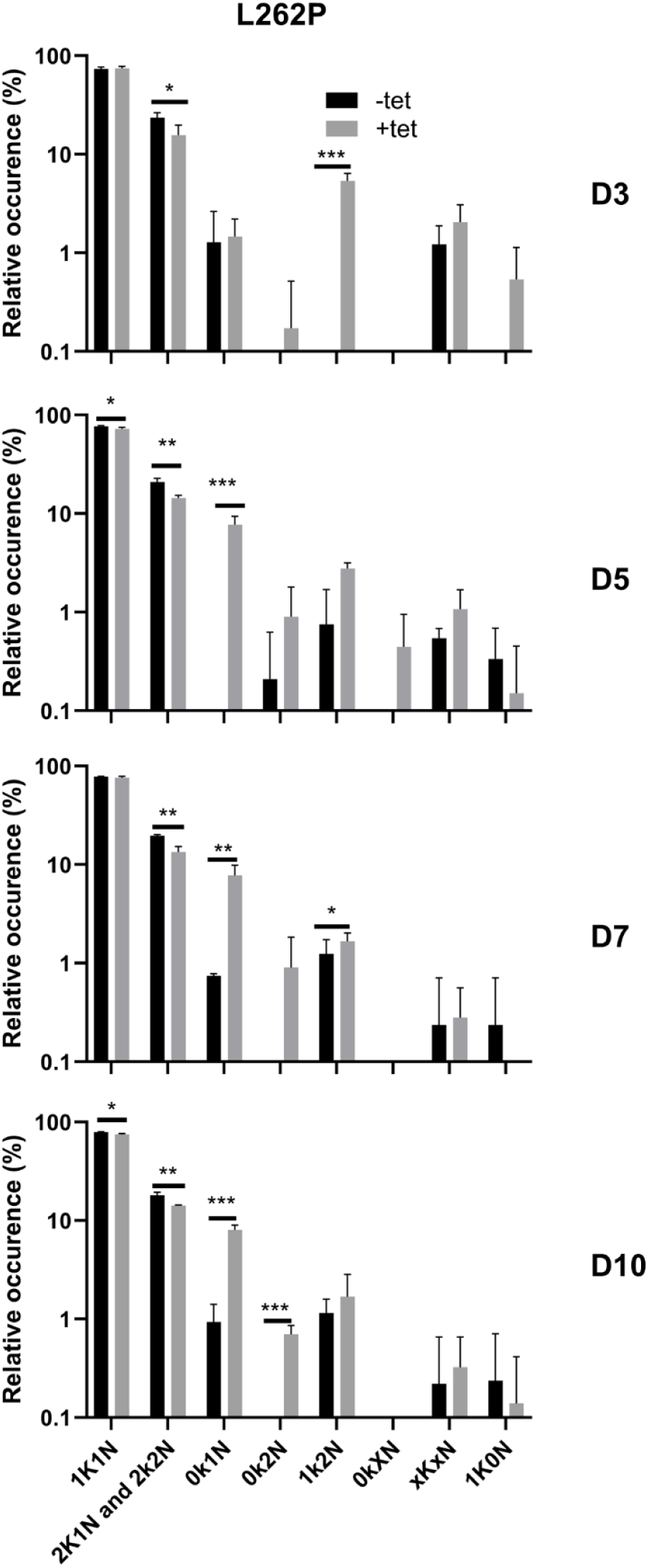
Quantification of relative occurrence of kDNA (K) and nuclei (N) in DAPI stained cells from induced (+tet) and non-induced (-tet) Tb927.8.4240 RNAi cells in the kDNA-independent γL262P background. Data for days 3, 5, 7 and 10 post-induction are shown (n>100 cells for each triplicate; unpaired t-test P<0.05*, <0.01**, <0.001***).

## REFERENCES

1. Stuart, K., Brun, R., Croft, S., Fairlamb, A., Gürtler, R.E., McKerrow, J., Reed, S. and Tarleton, R. (2008) Kinetoplastids: related protozoan pathogens, different diseases. The Journal of Clinical Investigation, 118, 1301–10.

2. Jensen, R.E. and Englund, P.T. (2012) Network News: The Replication of Kinetoplast DNA. Annual Review of Microbiology, 66, 473–491.

3. Kay, C., Williams, T.A. and Gibson, W. (2020) Mitochondrial DNAs provide insight into trypanosome phylogeny and molecular evolution. BMC Evolutionary Biology, 20, 161.

4. Alfonzo, J.D., Thiemann, O. and Simpson, L. (1997) The mechanism of U insertion/deletion RNA editing in kinetoplastid mitochondria. Nucleic acids research, 25, 3751–9.

5. Schnaufer, A., Domingo, G.J. and Stuart, K. (2002) Natural and induced dyskinetoplastic trypanosomatids: how to live without mitochondrial DNA. International Journal for Parasitology, 32, 1071–1084.

6. Lukes, J., Hashimi, H. and Zíková, A. (2005) Unexplained complexity of the mitochondrial genome and transcriptome in kinetoplastid flagellates. Current genetics, 48, 277–99.

7. Ogbadoyi, E.O., Robinson, D.R. and Gull, K. (2003) A High-Order Trans -Membrane Structural Linkage Is Responsible for Mitochondrial Genome Positioning and Segregation by Flagellar Basal Bodies in Trypanosomes. Molecular Biology of the Cell, 14, 1769–1779.

8. Englund, P.T. (2014) A passion for parasites. The Journal of biological chemistry, 289, 33712–29.

9. He, P., Katan, A.J., Tubiana, L., Dekker, C. and Michieletto, D. (2023) Single-Molecule Structure and Topology of Kinetoplast DNA Networks. Phys. Rev. X, 13, 021010.

10. Schneider, A. (2001) Unique aspects of mitochondrial biogenesis in trypanosomatids. International Journal for Parasitology, 31, 1403–1415.

11. Aphasizheva, I., Alfonzo, J., Carnes, J., Cestari, I., Cruz-Reyes, J., Göringer, H.U., Hajduk, S., Lukeš, J., Madison-Antenucci, S., Maslov, D.A., et al. (2020) Lexis and Grammar of Mitochondrial RNA Processing in Trypanosomes. Trends in Parasitology, 36, 337–355.

12. Cooper, S., Wadsworth, E.S., Ochsenreiter, T., Ivens, A., Savill, N.J. and Schnaufer, A. (2019) Assembly and annotation of the mitochondrial minicircle genome of a differentiation-competent strain of Trypanosoma brucei. Nucleic acids research, 47, 11304–11325.

13. Gould, M.K. and Schnaufer, A. (2014) Independence from Kinetoplast DNA Maintenance and Expression Is Associated with Multidrug Resistance in Trypanosoma brucei In Vitro. Antimicrobial Agents and Chemotherapy, 58, 2925–2928.

14. Roy Chowdhury, A., Bakshi, R., Wang, J., Yildirir, G., Liu, B., Pappas-Brown, V., Tolun, G., Griffith, J.D., Shapiro, T. a, Jensen, R.E., et al. (2010) The killing of African trypanosomes by ethidium bromide. PLoS Pathogens, 6, e1001226.

15. Schneider, A. and Ochsenreiter, T. (2018) Failure is not an option – mitochondrial genome segregation in trypanosomes. Journal of Cell Science, 131, jcs221820.

16. Amodeo, S., Bregy, I. and Ochsenreiter, T. (2022) Mitochondrial genome maintenance - the kinetoplast story. FEMS Microbiol Rev, 10.1093/femsre/fuac047.

17. Drew, M.E. and Englund, P.T. (2001) Intramitochondrial Location and Dynamics of Crithidia fasciculata Kinetoplast Minicircle Replication Intermediates. The Journal of Cell Biology, 153, 735–744.

18. Bruhn, D.F., Mozeleski, B., Falkin, L. and Klingbeil, M.M. (2010) Mitochondrial DNA polymerase POLIB is essential for minicircle DNA replication in African trypanosomes. Molecular Microbiology, 75, 1414–25.

19. Milman, N., Motyka, S.A., Englund, P.T., Robinson, D. and Shlomai, J. (2007) Mitochondrial origin-binding protein UMSBP mediates DNA replication and segregation in trypanosomes. Proceedings of the National Academy of Sciences of the United States of America, 104, 19250–5.

20. Wang, Z. and Englund, P.T. (2001) RNA interference of a trypanosome topoisomerase II causes progressive loss of mitochondrial DNA. The EMBO journal, 20, 4674–83.

21. Amodeo, S., Jakob, M. and Ochsenreiter, T. (2018) Characterization of the novel mitochondrial genome replication factor MiRF172 in Trypanosoma brucei. Journal of cell science, 131, pii: jcs211730.

22. Hines, J.C. and Ray, D.S. (2011) A second mitochondrial DNA primase is essential for cell growth and kinetoplast minicircle DNA replication in Trypanosoma brucei. Eukaryotic Cell, 10, 445–54.

23. Liu, B., Wang, J., Yaffe, N., Lindsay, M.E., Zhao, Z., Zick, A., Shlomai, J. and Englund, P.T. (2009) Trypanosomes have six mitochondrial DNA helicases with one controlling kinetoplast maxicircle replication. Molecular Cell, 35, 490–501.

24. Liu, B., Wang, J., Yildirir, G. and Englund, P.T. (2009) TbPIF5 is a Trypanosoma brucei mitochondrial DNA helicase involved in processing of minicircle Okazaki fragments. PLoS.Pathog., 5, e1000589.

25. Saxowsky, T.T., Choudhary, G., Klingbeil, M.M. and Englund, P.T. (2003) Trypanosoma brucei Has Two Distinct Mitochondrial DNA Polymerase β Enzymes. Journal of Biological Chemistry, 278, 49095–49101.

26. Downey, N., Hines, J.C., Sinha, K.M. and Ray, D.S. (2005) Mitochondrial DNA ligases of Trypanosoma brucei. Eukaryot Cell, 4, 765–774.

27. Amodeo, S., Bregy, I. and Ochsenreiter, T. (2023) Mitochondrial genome maintenance-the kinetoplast story. FEMS Microbiol Rev, 47, fuac047.

28. Carpenter, L.R. and Englund, P.T. (1995) Kinetoplast maxicircle DNA replication in Crithidia fasciculata and Trypanosoma brucei. Molecular and Cellular Biology, 15, 6794–6803.

29. Klingbeil, M.M., Motyka, S.A. and Englund, P.T. (2002) Multiple mitochondrial DNA polymerases in Trypanosoma brucei. Mol Cell, 10, 175–186.

30. Chandler, J., Vandoros, A.V., Mozeleski, B. and Klingbeil, M.M. (2008) Stem-loop silencing reveals that a third mitochondrial DNA polymerase, POLID, is required for kinetoplast DNA replication in trypanosomes. Eukaryot Cell, 7, 2141–2146.

31. Aeschlimann, S., Kalichava, A., Schimanski, B., Berger, B.M., Jetishi, C., Stettler, P., Ochsenreiter, T. and Schneider, A. (2022) Single p197 molecules of the mitochondrial genome segregation system of Trypanosoma brucei determine the distance between basal body and outer membrane. Proceedings of the National Academy of Sciences, 119, e2204294119.

32. Jetishi, C., Aeschlimann, S., Schimanski, B., Käser, S., Mullner, R., Oeljeklaus, S., Akiyoshi, B., Warscheid, B., Butter, F., Schneider, A., et al. (2025) Connecting basal body and mitochondrial DNA: TAC53 and the tubular organization of the tripartite attachment complex. PLoS Pathog, 21, e1013521.

33. Schimanski, B., Aeschlimann, S., Stettler, P., Käser, S., Gomez-Fabra Gala, M., Bender, J., Warscheid, B., Vögtle, F.-N. and Schneider, A. (2022) p166 links membrane and intramitochondrial modules of the trypanosomal tripartite attachment complex. PLOS Pathogens, 18, e1010207.

34. Baudouin, H.C.M., Pfeiffer, L. and Ochsenreiter, T. (2020) A comparison of three approaches for the discovery of novel tripartite attachment complex proteins in Trypanosoma brucei. PLOS Neglected Tropical Diseases, 14, e0008568.

35. Amodeo, S., Bregy, I., Hoffmann, A., Fradera-Sola, A., Kern, M., Baudouin, H., Zuber, B., Butter, F. and Ochsenreiter, T. (2023) Characterization of two novel proteins involved in mitochondrial DNA anchoring in Trypanosoma brucei. PLoS Pathog, 19, e1011486.

36. Pyrih, J., Hammond, M., Alves, A., Dean, S., Sunter, J.D., Wheeler, R.J., Gull, K. and Lukeš, J. (2023) Comprehensive sub-mitochondrial protein map of the parasitic protist Trypanosoma brucei defines critical features of organellar biology. Cell Rep, 42, 113083.

37. Horn, D. (2022) Genome-scale RNAi screens in African trypanosomes. Trends in parasitology, 38, 160–173.

38. Glover, L., Alsford, S., Baker, N., Turner, D.J., Sanchez-Flores, A., Hutchinson, S., Hertz-Fowler, C., Berriman, M. and Horn, D. (2015) Genome-scale RNAi screens for high-throughput phenotyping in bloodstream-form African trypanosomes. Nature protocols, 10, 106–33.

39. Hirumi, H. and Hirumi, K. (1989) Continuous cultivation of Trypanosoma brucei blood stream forms in a medium containing a low concentration of serum protein without feeder cell layers. The Journal of parasitology, 75, 985–9.

40. Dewar, C.E., MacGregor, P., Cooper, S., Gould, M.K., Matthews, K.R., Savill, N.J. and Schnaufer, A. (2018) Mitochondrial DNA is critical for longevity and metabolism of transmission stage Trypanosoma brucei. PLoS pathogens, 14, e1007195.

41. Creek, D.J., Nijagal, B., Kim, D.-H., Rojas, F., Matthews, K.R. and Barrett, M.P. (2013) Metabolomics guides rational development of a simplified cell culture medium for drug screening against Trypanosoma brucei. Antimicrobial Agents and Chemotherapy, 57, 2768–79.

42. Langmead, B. and Salzberg, S.L. (2012) Fast gapped-read alignment with Bowtie 2. Nature Methods, 9, 357–359.

43. Li, H., Handsaker, B., Wysoker, A., Fennell, T., Ruan, J., Homer, N., Marth, G., Abecasis, G. and Durbin, R. (2009) The Sequence Alignment/Map format and SAMtools. Bioinformatics, 25, 2078–2079.

44. Quinlan, A.R. (2014) BEDTools: The Swiss-Army Tool for Genome Feature Analysis. Curr Protoc Bioinformatics, 47, 11.12.1-34.

45. Robinson, M.D., McCarthy, D.J. and Smyth, G.K. (2010) edgeR: a Bioconductor package for differential expression analysis of digital gene expression data. Bioinformatics, 26, 139–140.

46. Inoue, M., Nakamura, Y., Yasuda, K., Yasaka, N., Hara, T., Schnaufer, A., Stuart, K. and Fukuma, T. (2005) The 14-3-3 Proteins of Trypanosoma brucei Function in Motility, Cytokinesis, and Cell Cycle. Journal of Biological Chemistry, 280, 14085–14096.

47. Burkard, G., Fragoso, C.M. and Roditi, I. (2007) Highly efficient stable transformation of bloodstream forms of Trypanosoma brucei. Molecular and Biochemical Parasitology, 153, 220–223.

48. Wirtz, E., Leal, S., Ochatt, C. and Cross, GeorgeA.M. (1999) A tightly regulated inducible expression system for conditional gene knock-outs and dominant-negative genetics in Trypanosoma brucei. Molecular and Biochemical Parasitology, 99, 89–101.

49. Livak, K.J. and Schmittgen, T.D. (2001) Analysis of relative gene expression data using real-time quantitative PCR and the 2(-Delta Delta C(T)) Method. Methods, 25, 402–408.

50. Brenndörfer, M. and Boshart, M. (2010) Selection of reference genes for mRNA quantification in Trypanosoma brucei. Molecular and biochemical parasitology, 172, 52–5.

51. Arena, E.T., Rueden, C.T., Hiner, M.C., Wang, S., Yuan, M. and Eliceiri, K.W. (2017) Quantitating the cell: turning images into numbers with ImageJ. Wiley Interdisciplinary Reviews: Developmental Biology, 6, e260.

52. Trikin, R., Doiron, N., Hoffmann, A., Haenni, B., Jakob, M., Schnaufer, A., Schimanski, B., Zuber, B. and Ochsenreiter, T. (2016) TAC102 Is a Novel Component of the Mitochondrial Genome Segregation Machinery in Trypanosomes. PLOS Pathogens, 12, e1005586.

53. Dean, S., Gould, M.K., Dewar, C.E. and Schnaufer, A.C. (2013) Single point mutations in ATP synthase compensate for mitochondrial genome loss in trypanosomes. Proceedings of the National Academy of Sciences, 110, 14741–14746.

54. Morris, J.C., Wang, Z., Drew, M.E. and Englund, P.T. (2002) Glycolysis modulates trypanosome glycoprotein expression as revealed by an RNAi library. The EMBO Journal, 21, 4429–4438.

55. Alsford, S., Turner, D.J., Obado, S.O., Sanchez-Flores, A., Glover, L., Berriman, M., Hertz-Fowler, C. and Horn, D. (2011) High-throughput phenotyping using parallel sequencing of RNA interference targets in the African trypanosome. Genome research, 21, 915–24.

56. Wang, Z., Drew, M.E., Morris, J.C. and Englund, P.T. (2002) Asymmetrical division of the kinetoplast DNA network of the trypanosome. The EMBO journal, 21, 4998–5005.

57. Ashley, N., Harris, D. and Poulton, J. (2005) Detection of mitochondrial DNA depletion in living human cells using PicoGreen staining. Experimental cell research, 303, 432–46.

58. Amodeo, S., Kalichava, A., Fradera-Sola, A., Bertiaux-Lequoy, E., Guichard, P., Butter, F. and Ochsenreiter, T. (2021) Characterization of the novel mitochondrial genome segregation factor TAP110 in Trypanosoma brucei. Journal of cell science, 134, jcs.254300.

59. Alvarez-Jarreta, J., Amos, B., Aurrecoechea, C., Bah, S., Barba, M., Barreto, A., Basenko, E.Y., Belnap, R., Blevins, A., Böhme, U., et al. (2024) VEuPathDB: the eukaryotic pathogen, vector and host bioinformatics resource center in 2023. Nucleic Acids Res, 52, D808–D816.

60. Billington, K., Halliday, C., Madden, R., Dyer, P., Barker, A.R., Moreira-Leite, F.F., Carrington, M., Vaughan, S., Hertz-Fowler, C., Dean, S., et al. (2023) Genome-wide subcellular protein map for the flagellate parasite Trypanosoma brucei. Nat Microbiol, 8, 533–547.

61. Zíková, A., Verner, Z., Nenarokova, A., Michels, P.A.M. and Lukeš, J. (2017) A paradigm shift: The mitoproteomes of procyclic and bloodstream Trypanosoma brucei are comparably complex. PLoS pathogens, 13, e1006679.

62. Panigrahi, A.K., Ogata, Y., Zíková, A., Anupama, A., Dalley, R.A., Acestor, N., Myler, P.J. and Stuart, K.D. (2009) A comprehensive analysis of Trypanosoma brucei mitochondrial proteome. Proteomics, 9, 434–50.

63. Peikert, C.D., Mani, J., Morgenstern, M., Käser, S., Knapp, B., Wenger, C., Harsman, A., Oeljeklaus, S., Schneider, A. and Warscheid, B. (2017) Charting organellar importomes by quantitative mass spectrometry. Nature Communications, 8, 15272.

64. Käser, S., Willemin, M., Schnarwiler, F., Schimanski, B., Poveda-Huertes, D., Oeljeklaus, S., Haenni, B., Zuber, B., Warscheid, B., Meisinger, C., et al. (2017) Biogenesis of the mitochondrial DNA inheritance machinery in the mitochondrial outer membrane of Trypanosoma brucei. PLOS Pathogens, 13, e1006808.

65. Käser, S., Oeljeklaus, S., Týč, J., Vaughan, S., Warscheid, B. and Schneider, A. (2016) Outer membrane protein functions as integrator of protein import and DNA inheritance in mitochondria. Proceedings of the National Academy of Sciences of the United States of America, 113, E4467–75.

66. Zhao, Z., Lindsay, M.E., Roy Chowdhury, A., Robinson, D.R. and Englund, P.T. (2008) p166, a link between the trypanosome mitochondrial DNA and flagellum, mediates genome segregation. The EMBO Journal, 27, 143–54.

67. Pyrih, J., Rašková, V., Škodová-Sveráková, I., Pánek, T. and Lukeš, J. (2020) ZapE/Afg1 interacts with Oxa1 and its depletion causes a multifaceted phenotype. PLOS ONE, 15, e0234918.

68. Nam, Y., Na, J., Ma, S.-X., Park, H., Park, H., Lee, E., Kim, H., Jang, S.-M., Ko, H.S. and Kim, S. (2024) DJ-1 protects cell death from a mitochondrial oxidative stress due to GBA1 deficiency. Genes Genomics, 10.1007/s13258-024-01506-w.

69. Benz, C. and Urbaniak, M.D. (2019) Organising the cell cycle in the absence of transcriptional control: Dynamic phosphorylation co-ordinates the Trypanosoma brucei cell cycle post-transcriptionally. PLoS Pathog, 15, e1008129.

70. Müller, M. and Papadopoulou, B. (2010) Stage-specific expression of the glycine cleavage complex subunits in Leishmania infantum. Mol Biochem Parasitol, 170, 17–27.

71. Sykes, S.E. and Hajduk, S.L. (2013) Dual functions of α-ketoglutarate dehydrogenase E2 in the Krebs cycle and mitochondrial DNA inheritance in Trypanosoma brucei. Eukaryotic Cell, 12, 78–90.

72. Gauba, E., Chen, H., Guo, L. and Du, H. (2019) Cyclophilin D deficiency attenuates mitochondrial F1Fo ATP synthase dysfunction via OSCP in Alzheimer’s disease. Neurobiol Dis, 121, 138–147.

73. Jackson, A.P., Allison, H.C., Barry, J.D., Field, M.C., Hertz-Fowler, C. and Berriman, M. (2013) A cell-surface phylome for African trypanosomes. PLoS Negl Trop Dis, 7, e2121.

74. Tonini, M.L., Peña-Diaz, P., Haindrich, A.C., Basu, S., Kriegová, E., Pierik, A.J., Lill, R., MacNeill, S.A., Smith, T.K. and Lukeš, J. (2018) Branched late-steps of the cytosolic iron-sulphur cluster assembly machinery of Trypanosoma brucei. PLoS Pathog, 14, e1007326.

75. Blum, M., Andreeva, A., Florentino, L.C., Chuguransky, S.R., Grego, T., Hobbs, E., Pinto, B.L., Orr, A., Paysan-Lafosse, T., Ponamareva, I., et al. (2025) InterPro: the protein sequence classification resource in 2025. Nucleic Acids Res, 53, D444–D456.

76. Jumper, J., Evans, R., Pritzel, A., Green, T., Figurnov, M., Ronneberger, O., Tunyasuvunakool, K., Bates, R., Žídek, A., Potapenko, A., et al. (2021) Highly accurate protein structure prediction with AlphaFold. Nature, 596, 583–589.

77. Shanmugasundram, A., Starns, D., Böhme, U., Amos, B., Wilkinson, P.A., Harb, O.S., Warrenfeltz, S., Kissinger, J.C., McDowell, M.A., Roos, D.S., et al. (2023) TriTrypDB: An integrated functional genomics resource for kinetoplastida. PLoS Negl Trop Dis, 17, e0011058.

78. Gluenz, E., Shaw, M.K. and Gull, K. (2007) Structural asymmetry and discrete nucleic acid subdomains in the Trypanosoma brucei kinetoplast. Molecular microbiology, 64, 1529–39.

79. Cadena, L.R., Svobodová, M., Benz, C., Rašková, V., Chmelová, Ľ., Škodová-Sveráková, I., Yurchenko, V., Lukeš, J., Hammond, M. and Durante, I.M. (2024) Characterization of novel and essential kinetoplast-associated proteins in Trypanosoma brucei. 10.1101/2024.04.22.590512.

80. Timms, M.W., van Deursen, F.J., Hendriks, E.F. and Matthews, K.R. (2002) Mitochondrial development during life cycle differentiation of African trypanosomes: evidence for a kinetoplast-dependent differentiation control point. Molecular Biology of the Cell, 13, 3747–59.

81. Bruhn, D.F., Sammartino, M.P. and Klingbeil, M.M. (2011) Three mitochondrial DNA polymerases are essential for kinetoplast DNA replication and survival of bloodstream form Trypanosoma brucei. Eukaryotic Cell, 10, 734–43.

82. Drozdz, M., Quijada, L. and Clayton, C.E. (2002) RNA interference in trypanosomes transfected with sense and antisense plasmids. Mol Biochem Parasitol, 121, 149–152.

83. Liu, Y. and Englund, P.T. (2007) The rotational dynamics of kinetoplast DNA replication. Mol Microbiol, 64, 676–690.

84. Wang, J., Englund, P.T. and Jensen, R.E. (2012) TbPIF8, a Trypanosoma brucei protein related to the yeast Pif1 helicase, is essential for cell viability and mitochondrial genome maintenance. Mol Microbiol, 83, 471–485.

85. Gluenz, E., Povelones, M.L., Englund, P.T. and Gull, K. (2011) The kinetoplast duplication cycle in Trypanosoma brucei is orchestrated by cytoskeleton-mediated cell morphogenesis. Molecular and cellular biology, 31, 1012–21.

86. Peña-Diaz, P., Vancová, M., Resl, C., Field, M.C. and Lukeš, J. (2017) A leucine aminopeptidase is involved in kinetoplast DNA segregation in Trypanosoma brucei. PLoS pathogens, 13, e1006310.

87. Cadena, L.R., Hammond, M., Tesařová, M., Chmelová, Ľ., Svobodová, M., Durante, I.M., Yurchenko, V. and Lukeš, J. (2024) A novel nabelschnur protein regulates segregation of the kinetoplast DNA in Trypanosoma brucei. Curr Biol, 34, 4803–4812.

88. Concepción-Acevedo, J., Luo, J. and Klingbeil, M.M. (2012) Dynamic Localization of Trypanosoma brucei Mitochondrial DNA Polymerase ID. Eukaryotic Cell, 11, 844–855.

89. Concepción-Acevedo, J., Miller, J.C., Boucher, M.J. and Klingbeil, M.M. (2018) Cell cycle localization dynamics of mitochondrial DNA polymerase IC in African trypanosomes. Molecular Biology of the Cell, 29, 2540–2552.

90. Sinha, K.M., Hines, J.C. and Ray, D.S. (2006) Cell cycle-dependent localization and properties of a second mitochondrial DNA ligase in Crithidia fasciculata. Eukaryot Cell, 5, 54–61.

91. Holehouse, A.S. and Kragelund, B.B. (2024) The molecular basis for cellular function of intrinsically disordered protein regions. Nat Rev Mol Cell Biol, 25, 187–211.

92. Lukes, J., Guilbride, D.L., Votýpka, J., Zíková, A., Benne, R. and Englund, P.T. (2002) Kinetoplast DNA network: evolution of an improbable structure. Eukaryotic cell, 1, 495–502.

93. Blom, D., De Haan, A., Van den Burg, J., van den Berg, M., Sloof, P., Jirku, M., Lukes, J. and Benne, R. (2000) Mitochondrial minicircles in the free-living bodonid Bodo saltans contain two gRNA gene cassettes and are not found in large networks. RNA, 6, 121–135.

94. Borst, P. (1991) Why kinetoplast DNA networks? Trends in genetics : TIG, 7, 139–41.

95. Kostygov, A.Y., Albanaz, A.T.S., Butenko, A., Gerasimov, E.S., Lukeš, J. and Yurchenko, V. (2023) Phylogenetic framework to explore trait evolution in Trypanosomatidae. Trends Parasitol, 40, 96–99.

96. Gaziová, I. and Lukes, J. (2003) Mitochondrial and nuclear localization of topoisomerase II in the flagellate Bodo saltans (Kinetoplastida), a species with non-catenated kinetoplast DNA. J Biol Chem, 278, 10900–10907.

97. Merkwirth, C. and Langer, T. (2009) Prohibitin function within mitochondria: Essential roles for cell proliferation and cristae morphogenesis. Biochimica et Biophysica Acta (BBA) - Molecular Cell Research, 1793, 27–32.

98. Edgar, R.C. (2004) MUSCLE: multiple sequence alignment with high accuracy and high throughput. Nucleic Acids Res, 32, 1792–1797.

99. Waterhouse, A.M., Procter, J.B., Martin, D.M.A., Clamp, M. and Barton, G.J. (2009) Jalview Version 2--a multiple sequence alignment editor and analysis workbench. *Bioinformatics*, **25**, 1189–1191.

100. Livingstone, C.D. and Barton, G.J. (1993) Protein sequence alignments: a strategy for the hierarchical analysis of residue conservation. Comput Appl Biosci, 9, 745–756.

