## Supplemental Table S1 for "A Genome-Wide Genetic Screen Identifies a Novel kDNA Replication Protein in Trypanosomes"

RNAi inserts were PCR-amplified from the transfected cell cultures after selection (see Materials and Methods) and before RNAi induction and FACS ("day 0 (D0)" samples; 5 replicates). The PCR amplicons were Illumina sequenced and reads mapped to the *T. brucei* TREU927 reference genome (release 42, v5.1; see Materials and Methods).

| | | **D0-1** | | **D0-2** | | **D0-3** | | **D0-4** | | **D0-5** | |
|---|---|---|---|---|---|---|---|---|---|---|---|
| **Contig** | **Reference length (bp)** | **Mapped reads** | **Fraction covered** | **Mapped reads** | **Fraction covered** | **Mapped reads** | **Fraction covered** | **Mapped reads** | **Fraction covered** | **Mapped reads** | **Fraction covered** |
| Tb927_01_v5.1 | 1,064,672 | 1,498,399 | 0.73 | 1,636,375 | 0.80 | 1,868,980 | 0.78 | 1,564,386 | 0.82 | 1,670,759 | 0.83 |
| Tb927_02_v5.1 | 1,193,948 | 1,534,941 | 0.76 | 1,834,204 | 0.83 | 1,457,321 | 0.81 | 1,347,609 | 0.84 | 1,658,320 | 0.86 |
| Tb927_03_v5.1 | 1,653,225 | 1,829,634 | 0.65 | 2,159,631 | 0.75 | 2,060,152 | 0.72 | 1,709,913 | 0.76 | 1,861,472 | 0.78 |
| Tb927_04_v5.1 | 1,590,432 | 2,020,247 | 0.68 | 2,401,433 | 0.77 | 1,766,594 | 0.74 | 1,465,184 | 0.78 | 1,680,832 | 0.80 |
| Tb927_05_v5.1 | 1,802,303 | 1,631,834 | 0.62 | 2,079,752 | 0.72 | 1,729,583 | 0.69 | 1,450,353 | 0.73 | 1,601,038 | 0.75 |
| Tb927_06_v5.1 | 1,618,915 | 1,985,131 | 0.67 | 2,346,411 | 0.77 | 2,119,947 | 0.73 | 1,663,130 | 0.77 | 1,711,680 | 0.79 |
| Tb927_07_v5.1 | 2,205,233 | 2,444,674 | 0.66 | 2,846,463 | 0.76 | 2,513,513 | 0.73 | 2,021,762 | 0.76 | 2,330,484 | 0.78 |
| Tb927_08_v5.1 | 2,481,190 | 2,159,880 | 0.67 | 3,030,098 | 0.76 | 2,370,025 | 0.74 | 2,245,222 | 0.77 | 2,435,429 | 0.79 |
| Tb927_09_v5.1 | 3,542,885 | 4,812,275 | 0.77 | 6,313,363 | 0.85 | 5,078,343 | 0.82 | 4,628,184 | 0.86 | 5,279,258 | 0.87 |
| Tb927_10_v5.1 | 4,144,375 | 4,015,510 | 0.63 | 5,071,283 | 0.73 | 3,944,557 | 0.70 | 3,317,422 | 0.73 | 3,540,325 | 0.76 |
| Tb927_11_v5.1 | 5,223,313 | 4,861,830 | 0.64 | 6,328,972 | 0.74 | 4,736,194 | 0.71 | 4,118,843 | 0.74 | 4,705,296 | 0.76 |
| Tb927_11_bin_v5.1 | 5,598,354 | 8,388,042 | 0.60 | 7,978,010 | 0.68 | 11,531,214 | 0.66 | 14,234,524 | 0.69 | 15,811,170 | 0.71 |
| Tb927_11_Homologues_1_v5.1 | 1,952 | 490 | 0.75 | 1,291 | 0.87 | 980 | 0.82 | 1,134 | 0.87 | 1,678 | 0.87 |
| Tb927_11_Homologues_2_v5.1 | 1,546 | 680 | 0.44 | 427 | 0.50 | 419 | 0.63 | 244 | 0.65 | 489 | 0.67 |
| Tb927_11_Homologues_3_v5.1 | 20,408 | 5,346 | 0.56 | 7,875 | 0.66 | 6,030 | 0.63 | 7,224 | 0.69 | 10,349 | 0.70 |
| Tb927_11_LH_fork_v5.1 | 14,430 | 16,734 | 0.49 | 17,629 | 0.63 | 13,305 | 0.55 | 19,272 | 0.67 | 8,932 | 0.68 |
| Tb927_11_RH_fork_v5.1 | 704,210 | 636,011 | 0.55 | 615,341 | 0.64 | 553,880 | 0.62 | 518,843 | 0.67 | 619,903 | 0.69 |
