## Supplemental Table S2 for "A Genome-Wide Genetic Screen Identifies a Novel kDNA Replication Protein in Trypanosomes"

Gene with genetic evidence for essential function in kDNA maintenance
Gene present in top 20 list
Gene investigated in more detail in this study
Gene present in reference set of 63 kinetoplast-associated proteins (KAP; Pyrih et al., 2023) = 7 genes
Gene present in reference set of 1650 mitochondrial proteins (Pyrih et al., 2023), but not in KAP set = 25 genes

Curated GO components in TriTrypDB indicate mitochondrial function or localisation = 32 genes

**TriTrypDB Curated GO Components**: Curated Gene Ontology Components from TriTrypDB
**TriTrypDB Curated GO Component IDs**: Gene Ontology Component IDs from TriTrypDB
**MitoTag Location Consensus**: Sublocalisation (Pyrih et al., 2023)

**Rank (by FDR1):** Rank Based on False Discovery Rate from D5 (+tet) vs D5 (-tet) Comparison
**TriTrypDB Annotation:** Gene Functional Annotation from TriTrypDB
**TriTrypDB ID**: TriTrypDB Gene Identifier (https://tritrypdb.org)
**logFC**: Log2 Fold Change
**logCPM**: Log Counts Per Million
**F**: F-statistic

| Rank (by FDR1) | TriTrypDB Annotation | TriTrypDB ID | D5 (+tet) vs D5 (-tet) | | | | | D5 (+tet) vs D0 | | | | | TriTrypDB Curated GO Components | TriTrypDB Curated GO Component IDs | MitoTag Location Consensus; Sublocalisation (Pyrih et al., 2023) |
|---|---|---|---|---|---|---|---|---|---|---|---|---|---|---|---|
| | | | logFC (tet+/tet-) | logCPM (tet+/tet-) | F (tet+/tet-) | PValue (tet+/tet-) | FDR (tet+/tet-) | logFC (tet+/D0) | logCPM (tet+/D0) | F (tet+/D0) | PValue (tet+/D0) | FDR (tet+/D0) | | | |
| 1 | **Tripartite Attachment Complex Protein 60** | ***Tb927.7.1400*** | 5.05 | 5.28 | 53.78 | 4.55E-07 | 2.15E-04 | 4.72 | 5.28 | 48.99 | 8.97E-07 | 3.23E-05 | cytoplasmic side of mitochondrial outer membrane;mitochondrion;tripartite attachmen | GO:0032473;GO:0005739;GO:0120121 | ***mito_TAC*** |
| 2 | **primase 2** | ***Tb927.1.4010*** | 5.49 | 13.36 | 53.98 | 4.66E-07 | 2.15E-04 | 8.27 | 13.36 | 95.41 | 5.45E-09 | 1.59E-05 | cytoplasm;mitochondrion | GO:0005737;GO:0005739 | |
| 3 | cyclophilin type peptidyl-prolyl cis-trans isomerase, putative | Tb927.10.13220 | 4.48 | 11.52 | 51.63 | 6.44E-07 | 2.60E-04 | 7.33 | 11.52 | 105.91 | 2.29E-09 | 8.96E-06 | nucleoplasm | GO:0005654 | |
| 4 | **Tripartite attachment complex protein 65** | ***Tb927.5.830*** | 6.41 | 9.98 | 49.44 | 8.81E-07 | 3.02E-04 | 6.02 | 9.98 | 45.28 | 1.64E-06 | 3.29E-05 | mitochondrion;tripartite attachment complex | GO:0005739;GO:0120121 | ***mito_TAC*** |
| 5 | **mitochondrial DNA polymerase I protein B** | ***Tb927.11.4690*** | 4.82 | 6.86 | 47.60 | 1.15E-06 | 3.46E-04 | 4.00 | 6.86 | 35.74 | 8.10E-06 | 5.47E-05 | antipodal site;cytoplasm;mitochondrial matrix;mitochondrion | GO:0140525;GO:0005737;GO:0005759;GO:0005 | ***Mito_KPE; Matrix/IMM peripheral proteins*** |
| 6 | prostaglandin f synthase | Tb927.11.4700 | 4.91 | 7.02 | 42.51 | 2.54E-06 | 6.08E-04 | 6.79 | 7.02 | 68.66 | 7.59E-08 | 3.23E-05 | ciliary plasm;cytoplasm;nuclear lumen;nucleoplasm | GO:0097014;GO:0005737;GO:0031981;GO:0005 | |
| 7 | hypothetical protein, conserved | ***Tb927.8.4240*** | 3.79 | 7.32 | 38.53 | 4.94E-06 | 1.05E-03 | 3.63 | 7.32 | 35.90 | 7.88E-06 | 5.38E-05 | ciliary basal body;kinetoplast;mitochondrial matrix;mitochondrion | GO:0036064;GO:0020023;GO:0005759;GO:0005 | ***Mito_KPE; Matrix/IMM peripheral proteins*** |
| 8 | DJ-1 family protein, putative | ***Tb927.6.2200*** | 6.41 | 6.72 | 37.28 | 6.14E-06 | 1.20E-03 | 4.68 | 6.72 | 23.76 | 9.58E-05 | 2.74E-04 | ciliary plasm;cytoplasm;cytosol;nucleoplasm;nucleus | GO:0097014;GO:0005737;GO:0005829;GO:0005 | ***n/a*** |
| 9 | cytosolic iron-sulfur protein assembly 1 | Tb927.8.3860 | 5.33 | 4.65 | 33.32 | 1.27E-05 | 1.86E-03 | 2.95 | 4.65 | 13.03 | 1.78E-03 | 3.27E-03 | CIA complex;ciliary plasm;cytoplasm;cytosol;nucleoplasm | GO:0097361;GO:0097014;GO:0005737;GO:0005 | |
| 10 | hypothetical protein, conserved | ***Tb927.6.2190*** | 5.79 | 6.59 | 29.72 | 2.59E-05 | 3.09E-03 | 4.86 | 6.59 | 23.13 | 1.11E-04 | 3.08E-04 | antipodal site;kinetoplast;mitochondrial matrix;mitochondrion;transport vesicle | GO:0140525;GO:0020023;GO:0005759;GO:0005 | ***Matrix/IMM peripheral proteins*** |
| 11 | **Tripartite attachment complex protein 166 (p166)** | ***Tb927.11.3290*** | 4.20 | 5.40 | 25.81 | 5.98E-05 | 5.61E-03 | 2.25 | 5.40 | 8.81 | 7.68E-03 | 1.24E-02 | kinetoplast;mitochondrial inner membrane;mitochondrion;tripartite attachment comple | GO:0020023;GO:0005743;GO:0005739;GO:012( | ***mito_TAC*** |
| 12 | variant surface glycoprotein (VSG, pseudogene), putative, variant surface glycoprotein (V | Tb927.9.16940 | 2.26 | 6.80 | 24.18 | 8.37E-05 | 7.40E-03 | 2.55 | 6.80 | 30.09 | 2.29E-05 | 9.78E-05 | N/A | N/A | |
| 13 | prefoldin subunit, putative | Tb927.11.16040 | 2.91 | 4.86 | 20.61 | 2.00E-04 | 1.28E-02 | 2.29 | 4.86 | 13.51 | 1.50E-03 | 2.81E-03 | cytoplasm | GO:0005737 | |
| 14 | hypothetical protein, conserved | Tb11.v5.0467 | 2.22 | 6.38 | 18.31 | 3.71E-04 | 1.88E-02 | 2.56 | 6.38 | 23.74 | 9.38E-05 | 2.69E-04 | N/A | N/A | |
| 15 | hypothetical protein (pseudogene) | Tb927.9.610 | 7.89 | 3.54 | 18.22 | 3.87E-04 | 1.93E-02 | 2.96 | 3.54 | 4.51 | 4.67E-02 | 6.49E-02 | N/A | N/A | |
| 16 | **Tripartite attachment complex protein 102** | ***Tb927.7.2390*** | 2.29 | 6.17 | 17.65 | 4.50E-04 | 2.13E-02 | 3.09 | 6.17 | 30.21 | 2.32E-05 | 9.87E-05 | mitochondrial matrix;mitochondrial protein-containing complex;mitochondrion;tripartit | GO:0005759;GO:0098798;GO:0005739;GO:012( | ***Mito_Kinetoplast; Matrix/IMM peripheral proteins*** |
| 17 | hypothetical protein, conserved | Tb927.10.11290 | 8.14 | 2.90 | 16.39 | 6.45E-04 | 2.55E-02 | 3.15 | 2.90 | 4.67 | 4.31E-02 | 6.04E-02 | N/A | N/A | |
| 18 | pyridoxal phosphate containing glycine decarboxylase, putative | ***Tb927.7.1910*** | 3.37 | 6.93 | 15.36 | 8.71E-04 | 2.97E-02 | 2.40 | 6.93 | 8.51 | 8.62E-03 | 1.38E-02 | cytoplasm;glycine cleavage complex;kinetoplast;mitochondrial inner membrane;mitoch | GO:0005737;GO:0005960;GO:0020023;GO:0005 | ***IMM integral*** |
| 19 | kinetoplastid-specific dual specificity phosphatase, putative | Tb927.2.4870 | 2.86 | 6.07 | 15.07 | 9.51E-04 | 3.09E-02 | 3.78 | 6.07 | 24.44 | 8.16E-05 | 2.42E-04 | ciliary plasm;cytoplasm;nuclear lumen | GO:0097014;GO:0005737;GO:0031981 | |
| 20 | hypothetical protein, conserved | Tb11.v5.0205 | 2.61 | 5.40 | 14.90 | 1.00E-03 | 3.11E-02 | 3.62 | 5.40 | 26.25 | 5.43E-05 | 1.79E-04 | cytoplasm | GO:0005737 | |
| 21 | 2OG-Fe(II) oxygenase superfamily, putative | ***Tb927.3.950*** | 2.06 | 7.83 | 14.36 | 1.18E-03 | 3.28E-02 | 3.08 | 7.83 | 29.75 | 2.57E-05 | 1.06E-04 | mitochondrion;nucleus | GO:0005739;GO:0005634 | ***n/a*** |
| 22 | phosphoacetylglucosamine mutase | Tb927.8.980 | 2.27 | 7.43 | 14.24 | 1.22E-03 | 3.35E-02 | 2.53 | 7.43 | 17.39 | 4.86E-04 | 1.04E-03 | ciliary plasm;cytoplasm;cytosol;glycosome | GO:0097014;GO:0005737;GO:0005829;GO:002( | |
| 23 | variant surface glycoprotein (VSG, pseudogene), putative | Tb927.9.18130 | 3.26 | 6.44 | 14.12 | 1.27E-03 | 3.43E-02 | 3.13 | 6.44 | 13.15 | 1.72E-03 | 3.16E-03 | N/A | N/A | |
| 24 | QA-SNARE protein putative | Tb927.9.13030 | 3.16 | 5.06 | 14.11 | 1.27E-03 | 3.43E-02 | 1.97 | 5.06 | 6.07 | 2.31E-02 | 3.41E-02 | SNARE complex;endomembrane system;integral component of membrane;transport ve | GO:0031201;GO:0012505;GO:0016021;GO:003( | |
| 25 | EF-hand domain-containing protein | Tb927.9.12340 | 2.05 | 5.74 | 14.04 | 1.28E-03 | 3.43E-02 | 1.89 | 5.74 | 12.00 | 2.46E-03 | 4.39E-03 | bilobe structure;cell tip;ciliary basal body;cytoplasm;nucleus;protein phosphatase type 2 | GO:0120120;GO:0051286;GO:0036064;GO:0005 | |
| 26 | hypothetical protein, conserved | Tb11.v5.0895 | 2.11 | 5.76 | 13.95 | 1.33E-03 | 3.49E-02 | 1.86 | 5.76 | 11.02 | 3.46E-03 | 5.99E-03 | N/A | N/A | |
| 27 | hypothetical protein, conserved | Tb927.10.6130 | 2.53 | 5.41 | 13.95 | 1.33E-03 | 3.49E-02 | 3.64 | 5.41 | 26.39 | 5.26E-05 | 1.75E-04 | cytoplasm | GO:0005737 | |
| 28 | RNA-editing substrate binding complex protein RESC14 | ***Tb927.9.7260*** | 2.01 | 5.71 | 13.57 | 1.48E-03 | 3.66E-02 | 3.07 | 5.71 | 29.12 | 2.81E-05 | 1.12E-04 | mitochondrial mRNA editing complex;mitochondrion | GO:0031019;GO:0005739 | ***n/a*** |
| 29 | kinesin heavy chain, putative | Tb927.1.1350 | 1.90 | 6.83 | 13.56 | 1.51E-03 | 3.71E-02 | 1.92 | 6.83 | 13.84 | 1.38E-03 | 2.61E-03 | cytoplasm;kinesin complex;microtubule | GO:0005737;GO:0005871;GO:0005874 | |
| 30 | hypothetical protein | Tb927.2.2015 | 3.80 | 6.61 | 13.52 | 1.53E-03 | 3.71E-02 | 2.63 | 6.61 | 7.21 | 1.44E-02 | 2.21E-02 | N/A | N/A | |
| 31 | variant surface glycoprotein (VSG, atypical), putative | Tb927.9.17910 | 2.57 | 6.05 | 13.40 | 1.59E-03 | 3.74E-02 | 3.21 | 6.05 | 19.83 | 2.52E-04 | 6.00E-04 | integral component of membrane | GO:0016021 | |
| 32 | KREPA3 | ***Tb927.8.620*** | 1.79 | 7.91 | 13.30 | 1.63E-03 | 3.79E-02 | 3.61 | 7.91 | 46.55 | 1.35E-06 | 3.23E-05 | antipodal site;kinetoplast;mitochondrial mRNA editing complex;mitochondrial matrix;m | GO:0140525;GO:0020023;GO:0031019;GO:0005 | ***Matrix/IMM peripheral proteins*** |
| 33 | Succinyl-CoA:3-ketoacid coenzyme A transferase, mitochondrial | ***Tb927.11.2690*** | 2.06 | 5.68 | 13.02 | 1.78E-03 | 3.95E-02 | 1.56 | 5.68 | 7.70 | 1.18E-02 | 1.84E-02 | cytoplasm;mitochondrion | GO:0005737;GO:0005739 | ***consensus mitoproteome*** |
| 34 | hypothetical protein, conserved | Tb927.8.6320 | 2.99 | 3.84 | 12.63 | 2.01E-03 | 4.21E-02 | 1.94 | 3.84 | 5.78 | 2.61E-02 | 3.82E-02 | cytoplasm | GO:0005737 | |
| 35 | zinc finger CCCH domain containing protein 44 | Tb927.11.7890 | 1.85 | 6.30 | 11.81 | 2.66E-03 | 4.79E-02 | 2.44 | 6.30 | 19.72 | 2.60E-04 | 6.15E-04 | cytoplasm | GO:0005737 | |
| 36 | U-box domain containing protein, putative | Tb927.3.3560 | 2.72 | 5.92 | 11.23 | 3.23E-03 | 5.32E-02 | 1.93 | 5.92 | 6.03 | 2.35E-02 | 3.46E-02 | axoneme;ciliary basal body;cytoplasm | GO:0005930;GO:0036064;GO:0005737 | |
| 37 | hypothetical protein, conserved | Tb927.6.3720 | 2.39 | 6.88 | 10.97 | 3.53E-03 | 5.59E-02 | 2.97 | 6.88 | 16.11 | 7.00E-04 | 1.43E-03 | ciliary plasm;cytoplasm;nuclear lumen | GO:0097014;GO:0005737;GO:0031981 | |
| 38 | expression site-associated gene 5 (ESAG5) protein, putative | Tb927.4.810 | 1.75 | 6.86 | 10.78 | 3.77E-03 | 5.76E-02 | 2.77 | 6.86 | 25.18 | 6.89E-05 | 2.13E-04 | cytoplasm;transport vesicle | GO:0005737;GO:0030133 | |
| 39 | STE/STE11 serine/threonine-protein kinase, putative | Tb927.10.10350 | 2.54 | 5.95 | 10.64 | 3.96E-03 | 5.89E-02 | 2.47 | 5.95 | 10.14 | 4.73E-03 | 7.96E-03 | ciliary basal body;cytoplasm | GO:0036064;GO:0005737 | |
| 40 | variant surface glycoprotein (VSG), putative | Tb11.v5.0599 | 2.22 | 5.64 | 10.59 | 4.03E-03 | 5.96E-02 | 2.92 | 5.64 | 17.41 | 4.84E-04 | 1.04E-03 | N/A | N/A | |
| 41 | hypothetical protein, conserved | Tb927.5.1170 | 2.51 | 4.05 | 10.34 | 4.35E-03 | 6.14E-02 | 3.61 | 4.05 | 19.54 | 2.65E-04 | 6.26E-04 | ciliary plasm;cytoplasm;nuclear lumen;nucleoplasm | GO:0097014;GO:0005737;GO:0031981;GO:0005 | |
| 42 | F-box protein, putative | Tb927.8.1380 | 2.29 | 4.05 | 10.27 | 4.45E-03 | 6.19E-02 | 1.79 | 4.05 | 6.56 | 1.86E-02 | 2.81E-02 | N/A | N/A | |
| 43 | hypothetical protein, conserved | Tb927.7.790 | 1.91 | 6.05 | 10.19 | 4.65E-03 | 6.30E-02 | 2.85 | 6.05 | 21.30 | 1.74E-04 | 4.41E-04 | cytoplasm | GO:0005737 | |
| 44 | expression site- associated gene 3 (ESAG3) protein, putative, expression site associated g | Tb10.v4.0032 | 1.99 | 6.92 | 10.17 | 4.67E-03 | 6.32E-02 | 3.31 | 6.92 | 25.28 | 6.74E-05 | 2.10E-04 | N/A | N/A | |
| 45 | stomatin-like protein, putative | ***Tb927.5.520*** | 1.99 | 7.57 | 9.95 | 5.06E-03 | 6.59E-02 | 3.86 | 7.57 | 32.17 | 1.59E-05 | 7.73E-05 | integral component of membrane;mitochondrion | GO:0016021;GO:0005739 | ***IMS/Membrane peripheral proteins*** |
| 46 | variant surface glycoprotein (VSG, pseudogene), putative, chrX additional, unordered cor | Tb10.v4.0212 | 2.21 | 5.65 | 9.91 | 5.12E-03 | 6.64E-02 | 2.90 | 5.65 | 16.23 | 6.75E-04 | 1.38E-03 | N/A | N/A | |
| 47 | glycosyltransferase (GlcNAc), putative | Tb927.2.2380 | 3.83 | 7.22 | 9.90 | 5.14E-03 | 6.65E-02 | 2.59 | 7.22 | 5.05 | 3.63E-02 | 5.16E-02 | cytoplasm;transport vesicle | GO:0005737;GO:0030133 | |
| 48 | lysyl-tRNA synthetase, putative | Tb927.8.1600 | 3.61 | 5.36 | 9.70 | 5.53E-03 | 6.98E-02 | 2.37 | 5.36 | 4.69 | 4.28E-02 | 6.00E-02 | cytoplasm;transport vesicle | GO:0005737;GO:0030133 | |
| 49 | kinetoplast ribosomal PPR-repeat containing protein 2 | ***Tb927.1.2990*** | 1.91 | 8.21 | 9.66 | 5.62E-03 | 7.02E-02 | 3.36 | 8.21 | 26.73 | 4.89E-05 | 1.66E-04 | antipodal site;cytoplasm;mitochondrial matrix;mitochondrial small ribosomal subunit;m | GO:0140525;GO:0005737;GO:0005759;GO:0005 | ***Matrix/IMM peripheral proteins*** |
| 50 | C/D snoRNA | Tb927_08_v4.snoRN/ | 4.10 | 2.69 | 9.44 | 6.09E-03 | 7.30E-02 | 3.05 | 2.69 | 6.00 | 2.38E-02 | 3.50E-02 | N/A | N/A | |
| 51 | expression site-associated gene (ESAG) protein, putative, expression site-associated gen | Tb927.10.4370 | 1.94 | 6.88 | 9.27 | 6.49E-03 | 7.49E-02 | 3.50 | 6.88 | 26.63 | 4.99E-05 | 1.68E-04 | cytoplasm;integral component of membrane;transport vesicle | GO:0005737;GO:0016021;GO:0030133 | |
| 52 | hypothetical protein, conserved | Tb927.10.7130 | 2.06 | 6.87 | 9.00 | 7.15E-03 | 7.87E-02 | 2.39 | 6.87 | 11.85 | 2.62E-03 | 4.64E-03 | cortical cytoskeleton | GO:0030863 | |
| 53 | Dicer-like protein 2 | Tb927.3.1230 | 1.59 | 7.73 | 8.89 | 7.47E-03 | 8.07E-02 | 3.13 | 7.73 | 30.53 | 2.20E-05 | 9.53E-05 | nucleoplasm;nucleus | GO:0005654;GO:0005634 | |
| 54 | Cytoskeleton-associated protein 67 kDa | Tb927.10.5880 | 1.68 | 7.97 | 8.67 | 8.11E-03 | 8.42E-02 | 3.50 | 7.97 | 32.58 | 1.47E-05 | 7.34E-05 | cortical cytoskeleton;cytoskeleton | GO:0030863;GO:0005856 | |
| 55 | expression site-associated gene 2 (ESAG2) protein, putative | Tb11.v5.0676 | 1.92 | 3.82 | 8.58 | 8.29E-03 | 8.55E-02 | 2.03 | 3.82 | 9.56 | 5.75E-03 | 9.53E-03 | N/A | N/A | |
| 56 | variant surface glycoprotein (VSG, atypical), putative | Tb927.3.370 | 1.73 | 5.30 | 8.50 | 8.58E-03 | 8.69E-02 | 2.91 | 5.30 | 22.20 | 1.36E-04 | 3.62E-04 | integral component of membrane | GO:0016021 | |
| 57 | C/D snoRNA | Tb927_08_v4.snoRN/ | 3.89 | 2.62 | 8.50 | 8.63E-03 | 8.69E-02 | 2.85 | 2.62 | 5.24 | 3.33E-02 | 4.76E-02 | N/A | N/A | |
| 58 | variant surface glycoprotein (VSG, pseudogene), putative, variant surface glycoprotein, p | Tb927.5.5040 | 1.84 | 6.54 | 8.46 | 8.78E-03 | 8.71E-02 | 2.17 | 6.54 | 11.54 | 2.91E-03 | 5.10E-03 | N/A | N/A | |
| 59 | hypothetical protein | Tb927.1.4970 | 1.64 | 5.55 | 8.17 | 9.79E-03 | 9.19E-02 | 2.32 | 5.55 | 15.71 | 7.80E-04 | 1.57E-03 | mitochondrion;transport vesicle | GO:0005739;GO:0030133 | |
| 60 | expression site-associated gene (ESAG) protein, putative | Tb11.01.6241 | 1.60 | 5.00 | 8.08 | 1.01E-02 | 9.31E-02 | 3.31 | 5.00 | 30.50 | 2.11E-05 | 9.26E-05 | N/A | N/A | |
| 61 | C/D snoRNA | Tb927_08_v4.snoRN/ | 3.69 | 2.70 | 7.97 | 1.06E-02 | 9.52E-02 | 2.82 | 2.70 | 5.24 | 3.33E-02 | 4.76E-02 | N/A | N/A | |
| 62 | glycosyltransferase (GlcNAc), putative | Tb927.2.2400 | 1.63 | 6.46 | 7.94 | 1.07E-02 | 9.55E-02 | 2.36 | 6.46 | 15.89 | 7.46E-04 | 1.51E-03 | cytoplasm;mitochondrion;transport vesicle | GO:0005737;GO:0005739;GO:0030133 | |
| 63 | transcription factor IIa, putative | Tb927.10.4840 | 2.71 | 3.14 | 7.82 | 1.13E-02 | 9.78E-02 | 4.04 | 3.14 | 15.67 | 7.95E-04 | 1.59E-03 | cytoplasm;nuclear lumen | GO:0005737;GO:0031981 | |
| 64 | variant surface glycoprotein (VSG), putative | Tb11.v5.0923 | 1.74 | 6.34 | 7.74 | 1.16E-02 | 9.93E-02 | 1.99 | 6.34 | 10.02 | 4.93E-03 | 8.29E-03 | N/A | N/A | |
| 65 | hypothetical protein, conserved | Tb927.5.3080 | 2.34 | 5.94 | 7.73 | 1.17E-02 | 9.95E-02 | 1.81 | 5.94 | 4.80 | 4.06E-02 | 5.71E-02 | ciliary basal body;ciliary pro-basal body | GO:0036064;GO:0120280 | |
| 66 | hypothetical protein | Tb927.3.3600 | 1.81 | 5.13 | 7.67 | 1.19E-02 | 1.01E-01 | 2.35 | 5.13 | 12.42 | 2.17E-03 | 3.91E-03 | ciliary plasm;cytoplasm;nuclear lumen | GO:0097014;GO:0005737;GO:0031981 | |
| 67 | variant surface glycoprotein, degenerate, variant surface glycoprotein (VSG, pseudogene | Tb927.3.350 | 1.65 | 5.16 | 7.63 | 1.21E-02 | 1.01E-01 | 1.59 | 5.16 | 7.20 | 1.43E-02 | 2.21E-02 | N/A | N/A | |
| 68 | hypothetical protein, conserved | Tb927.4.4100 | 1.70 | 6.75 | 7.59 | 1.23E-02 | 1.02E-01 | 3.26 | 6.75 | 24.76 | 7.58E-05 | 2.29E-04 | cytoplasm;mitochondrion;nuclear lumen;ribonucleoprotein complex | GO:0005737;GO:0005739;GO:0031981;GO:199( | |
| 69 | expression site-associated gene (ESAG, pseudogene), putative, expression site-associate | Tb927.3.5800 | 2.28 | 4.98 | 7.39 | 1.34E-02 | 1.07E-01 | 4.00 | 4.98 | 19.68 | 2.63E-04 | 6.20E-04 | N/A | N/A | |
| 70 | NADH-ubiquinone oxidoreductase B22 subunit | ***Tb927.11.15810*** | 1.73 | 7.41 | 7.36 | 1.35E-02 | 1.07E-01 | 3.62 | 7.41 | 27.86 | 3.83E-05 | 1.38E-04 | NADH dehydrogenase complex;mitochondrial inner membrane;mitochondrial membran | GO:0030964;GO:0005743;GO:0031966;GO:0005 | ***IMM integral*** |
| 71 | variant surface glycoprotein (VSG, pseudogene), putative | Tb927.10.16320 | 3.51 | 4.96 | 7.34 | 1.36E-02 | 1.07E-01 | 2.66 | 4.96 | 4.57 | 4.53E-02 | 6.32E-02 | N/A | N/A | |
| 72 | hypothetical protein, conserved | Tb927.7.2280 | 2.68 | 6.01 | 7.30 | 1.38E-02 | 1.08E-01 | 3.22 | 6.01 | 10.12 | 4.76E-03 | 8.02E-03 | cytoplasm;mitochondrion;paraflagellar rod | GO:0005737;GO:0005739;GO:0097740 | |
| 73 | hypothetical protein, conserved | Tb927.11.14640 | 1.68 | 5.53 | 7.23 | 1.42E-02 | 1.09E-01 | 2.33 | 5.53 | 13.42 | 1.57E-03 | 2.92E-03 | cytoplasm | GO:0005737 | |
| 74 | hypothetical protein, conserved | ***Tb927.10.11340*** | 2.47 | 8.31 | 7.23 | 1.42E-02 | 1.09E-01 | 4.64 | 8.31 | 21.02 | 1.86E-04 | 4.67E-04 | cytoplasm | GO:0005737 | ***n/a*** |
| 75 | C/D snoRNA | Tb927_08_v4.snoRN/ | 3.69 | 2.70 | 7.11 | 1.50E-02 | 1.12E-01 | 2.86 | 2.70 | 4.76 | 4.15E-02 | 5.83E-02 | N/A | N/A | |
| 76 | FLA1-like protein | Tb927.10.6180 | 1.59 | 6.04 | 7.10 | 1.50E-02 | 1.12E-01 | 2.84 | 6.04 | 20.71 | 2.01E-04 | 4.98E-04 | acidocalcisome | GO:0020022 | |
| 77 | EF-hand domain pair, putative | Tb927.11.6460 | 1.62 | 8.01 | 7.07 | 1.52E-02 | 1.13E-01 | 3.44 | 8.01 | 27.89 | 3.79E-05 | 1.38E-04 | axoneme;cilium;cytoplasm | GO:0005930;GO:0005929;GO:0005737 | |
| 78 | protein kinase, putative | Tb927.3.3290 | 2.13 | 4.32 | 7.05 | 1.53E-02 | 1.14E-01 | 1.86 | 4.32 | 5.49 | 2.98E-02 | 4.30E-02 | cell tip;cytoplasm;transport vesicle | GO:0051286;GO:0005737;GO:0030133 | |
| 79 | C/D snoRNA | Tb927_08_v4.snoRN/ | 3.42 | 2.76 | 6.92 | 1.61E-02 | 1.17E-01 | 2.95 | 2.76 | 5.52 | 2.93E-02 | 4.24E-02 | N/A | N/A | |
| 80 | mitochondrial carrier protein, putative | ***Tb927.4.1660*** | 1.89 | 5.07 | 6.83 | 1.67E-02 | 1.18E-01 | 1.74 | 5.07 | 5.89 | 2.49E-02 | 3.66E-02 | glycosome;mitochondrial inner membrane;mitochondrial membrane;mitochondrion | GO:0020015;GO:0005743;GO:0031966;GO:0005 | ***IMM integral*** |
| 81 | hypothetical protein, conserved | Tb927.4.1550 | 1.51 | 6.25 | 6.76 | 1.73E-02 | 1.21E-01 | 1.78 | 6.25 | 9.18 | 6.69E-03 | 1.09E-02 | cytoplasm;nucleoplasm | GO:0005737;GO:0005654 | |

| | | | | | | | | | | | | | | | |
|---|---|---|---|---|---|---|---|---|---|---|---|---|---|---|---|
| 82 | protein kinase, putative | Tb11.v5.0474 | 2.09 | 4.27 | 6.71 | 1.76E-02 | 1.22E-01 | 1.80 | 4.27 | 5.09 | 3.55E-02 | 5.05E-02 | N/A | N/A | |
| 83 | NADH dehydrogenase (ubiquinone) 1 alpha subcomplex subunit, putative | ***Tb927.2.4380*** | 1.75 | 9.30 | 6.70 | 1.77E-02 | 1.22E-01 | 5.37 | 9.30 | 45.83 | 1.51E-06 | 3.24E-05 | NADH dehydrogenase complex;cytoplasm;kinetoplast;mitochondrial matrix;mitochondr | GO:0030964;GO:0005737;GO:0020023;GO:0005 | ***Matrix/IMM peripheral proteins*** |
| 84 | zinc finger CCCH domain containing protein 12 | Tb927.5.1570 | 2.07 | 7.22 | 6.66 | 1.80E-02 | 1.24E-01 | 2.78 | 7.22 | 11.37 | 3.08E-03 | 5.38E-03 | cytoplasm;cytosol;transport vesicle | GO:0005737;GO:0005829;GO:0030133 | |
| 85 | hypothetical protein, conserved | ***Tb927.10.5570*** | 1.93 | 4.26 | 6.62 | 1.82E-02 | 1.25E-01 | 2.97 | 4.26 | 14.49 | 1.12E-03 | 2.17E-03 | cytoplasm;kinetoplast;mitochondrial inner membrane;mitochondrion | GO:0005737;GO:0020023;GO:0005743;GO:0005 | ***IMM integral*** |
| 86 | Histone RNA hairpin-binding protein RNA-binding domain containing protein, putative | Tb927.3.870 | 1.73 | 5.02 | 6.57 | 1.87E-02 | 1.26E-01 | 2.94 | 5.02 | 17.35 | 4.92E-04 | 1.05E-03 | cytoplasm;histone pre-mRNA 3'end processing complex;nucleus | GO:0005737;GO:0071204;GO:0005634 | |
| 87 | variant surface glycoprotein (VSG, pseudogene), putative, chrIX additional, unordered co | Tb09.v4.0186 | 2.02 | 3.69 | 6.53 | 1.89E-02 | 1.26E-01 | 2.99 | 3.69 | 13.48 | 1.52E-03 | 2.84E-03 | N/A | N/A | |
| 88 | variant surface protein (VSG), putative | Tb11.v5.0138 | 1.98 | 4.49 | 6.54 | 1.89E-02 | 1.26E-01 | 2.97 | 4.49 | 13.68 | 1.45E-03 | 2.72E-03 | N/A | N/A | |
| 89 | 5.8S ribosomal RNA | Tb927.11.rRNA_1:rRI | 1.81 | 3.19 | 6.52 | 1.90E-02 | 1.26E-01 | 1.62 | 3.19 | 5.37 | 3.12E-02 | 4.49E-02 | N/A | N/A | |
| 90 | hypothetical protein, conserved | Tb927.10.4820 | 2.52 | 2.72 | 6.50 | 1.92E-02 | 1.27E-01 | 2.22 | 2.72 | 5.26 | 3.29E-02 | 4.71E-02 | bilobe structure | GO:0120120 | |
| 91 | hypothetical protein, conserved | ***Tb927.7.6410*** | 2.02 | 7.06 | 6.47 | 1.95E-02 | 1.28E-01 | 3.01 | 7.06 | 13.36 | 1.61E-03 | 2.98E-03 | mitochondrial matrix;mitochondrion | GO:0005759;GO:0005739 | ***Matrix/IMM peripheral proteins*** |
| 92 | hypothetical protein, conserved | Tb927.10.5530 | 1.56 | 6.03 | 6.41 | 2.00E-02 | 1.30E-01 | 1.93 | 6.03 | 9.58 | 5.79E-03 | 9.59E-03 | cytoplasm | GO:0005737 | |
| 93 | vacuolar protein sorting-associated protein 35, putative | Tb927.10.2270 | 1.65 | 11.62 | 6.40 | 2.01E-02 | 1.31E-01 | 3.94 | 11.62 | 30.51 | 2.21E-05 | 9.55E-05 | cytoplasm;late endosome;membrane;retromer complex;transport vesicle | GO:0005737;GO:0005770;GO:0016020;GO:0030 | |
| 94 | Calpain-like protein 1.1 | Tb927.1.2100 | 1.76 | 6.08 | 6.39 | 2.01E-02 | 1.31E-01 | 1.56 | 6.08 | 5.11 | 3.53E-02 | 5.02E-02 | ciliary tip;paraflagellar rod | GO:0097542;GO:0097740 | |
| 95 | Noncoding RNA, putative | Tb7.NT.53:ncRNA | 2.54 | -0.91 | 6.34 | 2.06E-02 | 1.33E-01 | 1.91 | -0.91 | 5.12 | 3.51E-02 | 5.01E-02 | N/A | N/A | |
| 96 | C/D snoRNA | Tb927_08_v4.snoRN/ | 3.16 | 2.75 | 6.27 | 2.12E-02 | 1.35E-01 | 2.84 | 2.75 | 5.33 | 3.19E-02 | 4.58E-02 | N/A | N/A | |
| 97 | hypothetical protein, conserved | ***Tb927.10.12160*** | 1.73 | 4.97 | 6.25 | 2.14E-02 | 1.35E-01 | 1.77 | 4.97 | 6.56 | 1.87E-02 | 2.82E-02 | ciliary plasm;cytoplasm;mitochondrial matrix;mitochondrion;nuclear lumen | GO:0097014;GO:0005737;GO:0005759;GO:0005 | ***Matrix/IMM peripheral proteins*** |
| 98 | Mechanosensitive ion channel MscS, putative | ***Tb927.10.9030*** | 2.40 | 2.66 | 5.96 | 2.43E-02 | 1.43E-01 | 2.66 | 2.66 | 7.28 | 1.40E-02 | 2.16E-02 | ciliary plasm;cytoplasm;kinetoplast;mitochondrial matrix;mitochondrion;nuclear lumen; | GO:0097014;GO:0005737;GO:0020023;GO:0005 | ***Matrix/IMM peripheral proteins*** |
| 99 | mitochondrial ATP synthase subunit ATPTB11 | ***Tb927.3.2180*** | 1.59 | 7.04 | 5.86 | 2.52E-02 | 1.46E-01 | 3.13 | 7.04 | 20.27 | 2.25E-04 | 5.47E-04 | mitochondrial inner membrane;mitochondrial proton-transporting ATP synthase comple | GO:0005743;GO:0005753;GO:0005739 | ***IMM integral*** |
| 100 | hypothetical protein, conserved | Tb927.11.15250 | 1.85 | 6.10 | 5.84 | 2.55E-02 | 1.47E-01 | 4.26 | 6.10 | 25.15 | 6.94E-05 | 2.14E-04 | axoneme;cilium | GO:0005930;GO:0005929 | |
| 101 | pteridine transporter, putative | Tb927.1.2820 | 1.61 | 6.80 | 5.83 | 2.56E-02 | 1.47E-01 | 2.56 | 6.80 | 13.81 | 1.40E-03 | 2.63E-03 | N/A | N/A | |
| 102 | variant surface glycoprotein (VSG, pseudogene), putative, chrIX additional, unordered co | Tb09.v4.0057 | 1.66 | 4.58 | 5.71 | 2.69E-02 | 1.51E-01 | 3.16 | 4.58 | 18.55 | 3.50E-04 | 7.89E-04 | N/A | N/A | |
| 103 | malate dehydrogenase-related | ***Tb927.10.2550*** | 1.61 | 6.46 | 5.68 | 2.73E-02 | 1.52E-01 | 2.79 | 6.46 | 15.68 | 7.93E-04 | 1.59E-03 | cytoplasm;mitochondrial matrix;mitochondrion | GO:0005737;GO:0005759;GO:0005739 | ***Matrix/IMM peripheral proteins*** |
| 104 | pteridine transporter, putative | ***Tb927.1.2880*** | 1.58 | 5.58 | 5.65 | 2.77E-02 | 1.53E-01 | 1.68 | 5.58 | 6.40 | 2.00E-02 | 3.00E-02 | N/A | N/A | ***n/a*** |
| 105 | expression site-associated gene 2 (ESAG2) protein, putative | Tb11.v5.0789 | 1.63 | 3.58 | 5.62 | 2.80E-02 | 1.54E-01 | 2.71 | 3.58 | 14.45 | 1.12E-03 | 2.16E-03 | N/A | N/A | |
| 106 | hypothetical protein, conserved | Tb927.3.890 | 1.58 | 5.38 | 5.59 | 2.85E-02 | 1.55E-01 | 2.86 | 5.38 | 16.65 | 6.00E-04 | 1.25E-03 | N/A | N/A | |
| 107 | variant surface glycoprotein (VSG, atypical), putative | Tb927.5.4730 | 1.98 | 4.38 | 5.53 | 2.92E-02 | 1.58E-01 | 1.94 | 4.38 | 5.37 | 3.13E-02 | 4.51E-02 | integral component of membrane | GO:0016021 | |
| 108 | class I transcription factor A, subunit 5b | Tb927.8.4130 | 1.58 | 4.25 | 5.52 | 2.92E-02 | 1.58E-01 | 1.82 | 4.25 | 7.18 | 1.44E-02 | 2.22E-02 | N/A | N/A | |
| 109 | pteridine transporter, putative | Tb927.1.2850 | 1.52 | 6.71 | 5.44 | 3.04E-02 | 1.61E-01 | 2.58 | 6.71 | 14.45 | 1.14E-03 | 2.20E-03 | N/A | N/A | |
| 110 | variant surface glycoprotein (VSG), degenerate, variant surface glycoprotein (VSG, pseud | Tb927.9.16170 | 1.63 | 6.94 | 5.43 | 3.05E-02 | 1.61E-01 | 3.19 | 6.94 | 18.56 | 3.53E-04 | 7.95E-04 | N/A | N/A | |
| 111 | SUMO-interacting motif-containing protein | ***Tb927.1.4410*** | 1.51 | 9.07 | 5.37 | 3.14E-02 | 1.63E-01 | 3.33 | 9.07 | 22.79 | 1.21E-04 | 3.29E-04 | cytoplasm;mitochondrion;nucleoplasm;transport vesicle | GO:0005737;GO:0005739;GO:0005654;GO:0030 | ***IMS/Membrane peripheral proteins*** |
| 112 | hypothetical protein, conserved | Tb927.7.3870 | 1.51 | 6.42 | 5.31 | 3.22E-02 | 1.66E-01 | 4.19 | 6.42 | 32.96 | 1.36E-05 | 7.07E-05 | N/A | N/A | |
| 113 | hypothetical protein, conserved | Tb927.11.12200 | 1.52 | 6.89 | 5.19 | 3.40E-02 | 1.70E-01 | 4.08 | 6.89 | 30.56 | 2.18E-05 | 9.48E-05 | endoplasmic reticulum;nuclear envelope | GO:0005783;GO:0005635 | |
| 114 | hypothetical protein, conserved | Tb927.2.5970 | 1.84 | 7.01 | 5.18 | 3.42E-02 | 1.71E-01 | 4.31 | 7.01 | 23.11 | 1.11E-04 | 3.09E-04 | ciliary pocket collar | GO:1990900 | |
| 115 | expression site-associated gene (ESAG, pseudogene), putative, chrIX additional, unorder | Tb09.v4.0154 | 1.55 | 6.49 | 5.17 | 3.44E-02 | 1.71E-01 | 3.54 | 6.49 | 23.11 | 1.11E-04 | 3.09E-04 | N/A | N/A | |
| 116 | hypothetical protein, conserved | Tb927.9.8620 | 1.52 | 9.64 | 5.09 | 3.56E-02 | 1.74E-01 | 4.53 | 9.64 | 35.01 | 9.27E-06 | 5.82E-05 | ciliary basal body;ciliary pocket collar | GO:0036064;GO:1990900 | |
| 117 | invariant surface glycoprotein, putative | Tb927.5.420 | 1.56 | 7.39 | 5.08 | 3.57E-02 | 1.75E-01 | 3.92 | 7.39 | 26.72 | 4.89E-05 | 1.66E-04 | transport vesicle | GO:0030133 | |
| 118 | syntaxin, putative | Tb927.10.1830 | 1.59 | 5.01 | 5.06 | 3.61E-02 | 1.76E-01 | 2.87 | 5.01 | 14.95 | 9.84E-04 | 1.93E-03 | SNARE complex;endomembrane system;integral component of membrane;membrane;p | GO:0031201;GO:0012505;GO:0016021;GO:0016 | |
| 119 | ABC transporter, putative | Tb927.9.6310 | 1.73 | 9.06 | 5.06 | 3.61E-02 | 1.76E-01 | 3.77 | 9.06 | 20.38 | 2.19E-04 | 5.33E-04 | cytoplasmic side of plasma membrane;endoplasmic reticulum;integral component of me | GO:0009898;GO:0005783;GO:0016021;GO:0005 | |
| 120 | maoC-like dehydratase, putative | ***Tb927.8.1440*** | 1.79 | 3.78 | 4.99 | 3.71E-02 | 1.79E-01 | 3.16 | 3.78 | 14.11 | 1.25E-03 | 2.38E-03 | ciliary plasm;cytoplasm;mitochondrial matrix;mitochondrion;nuclear lumen;tripartite att | GO:0097014;GO:0005737;GO:0005759;GO:0005 | ***Matrix/IMM peripheral proteins*** |
| 121 | DNA cross-link repair protein pso2/snm1 | Tb927.4.1480 | 1.77 | 6.65 | 4.96 | 3.79E-02 | 1.81E-01 | 2.06 | 6.65 | 6.61 | 1.84E-02 | 2.78E-02 | nuclear lumen;nucleus | GO:0031981;GO:0005634 | |
| 122 | cAMP response protein 10 | Tb927.11.7180 | 1.64 | 9.09 | 4.94 | 3.81E-02 | 1.82E-01 | 4.65 | 9.09 | 30.69 | 2.13E-05 | 9.31E-05 | axoneme | GO:0005930 | |
| 123 | Exocyst complex component EXO99 | Tb927.10.420 | 2.09 | 6.02 | 4.94 | 3.81E-02 | 1.82E-01 | 2.53 | 6.02 | 6.99 | 1.57E-02 | 2.40E-02 | ciliary pocket;exocyst | GO:0020016;GO:0000145 | |
| 124 | chrXI additional, unordered contigs, variant surface glycoprotein (VSG), putative | Tb11.0590 | 2.53 | 2.78 | 4.81 | 4.04E-02 | 1.89E-01 | 4.43 | 2.78 | 12.48 | 2.13E-03 | 3.84E-03 | N/A | N/A | |
| 125 | adenosine kinase, putative | Tb927.6.2360 | 1.55 | 8.70 | 4.74 | 4.18E-02 | 1.92E-01 | 3.80 | 8.70 | 23.72 | 9.66E-05 | 2.75E-04 | ciliary plasm;cytoplasm;cytosol;glycosome;nucleoplasm;nucleus | GO:0097014;GO:0005737;GO:0005829;GO:0020 | |
| 126 | variant surface glycoprotein (VSG, pseudogene), putative, chrIX additional, unordered co | Tb09.v4.0180 | 2.54 | 3.52 | 4.69 | 4.29E-02 | 1.95E-01 | 2.86 | 3.52 | 5.81 | 2.58E-02 | 3.78E-02 | N/A | N/A | |
| 127 | hypothetical protein, conserved | Tb927.8.5270 | 2.10 | 5.37 | 4.60 | 4.47E-02 | 2.00E-01 | 3.81 | 5.37 | 13.05 | 1.77E-03 | 3.25E-03 | N/A | N/A | |
| 128 | hypothetical protein, conserved | Tb927.7.1650 | 1.63 | 8.01 | 4.55 | 4.57E-02 | 2.02E-01 | 3.83 | 8.01 | 21.18 | 1.79E-04 | 4.52E-04 | nuclear lumen;nucleoplasm;nucleus | GO:0031981;GO:0005654;GO:0005634 | |
| 129 | RNA-editing substrate binding complex protein RESC15 | ***Tb927.1.1730*** | 1.50 | 5.47 | 4.50 | 4.67E-02 | 2.05E-01 | 1.93 | 5.47 | 7.23 | 1.43E-02 | 2.20E-02 | mitochondrial mRNA editing complex;mitochondrial matrix;mitochondrion | GO:0031019;GO:0005759;GO:0005739 | ***Matrix/IMM peripheral proteins*** |
| 130 | variant surface glycoprotein, fragment | Tb927.10.16300 | 1.79 | 3.58 | 4.48 | 4.71E-02 | 2.05E-01 | 3.41 | 3.58 | 14.41 | 1.14E-03 | 2.20E-03 | N/A | N/A | |
| 131 | pre-mRNA processing factor 4 (PRP4) like/WD domain, G-beta repeat, putative | Tb11.v5.0631 | 1.69 | 6.31 | 4.43 | 4.84E-02 | 2.05E-01 | 2.86 | 6.31 | 11.67 | 2.78E-03 | 4.90E-03 | U4/U6 x U5 tri-snRNP complex;cytoplasm;nucleus | GO:0046540;GO:0005737;GO:0005634 | |
| 132 | hypothetical protein, conserved | ***Tb927.11.16530*** | 2.02 | 6.31 | 4.37 | 4.97E-02 | 2.07E-01 | 2.63 | 6.31 | 7.12 | 1.49E-02 | 2.28E-02 | mitochondrion | GO:0005739 | ***n/a*** |
