## Supplemental Table S2 for "A Genome-Wide Genetic Screen Identifies a Novel kDNA Replication Protein in Trypanosomes"

| Primer Name | Primer direction | Primer sequence 5’-3’ | Description |
|---|---|---|---|
| LIB2f | F | TAGCCCCTCGAGGGCCAGT | RNAi inserts amplification |
| LIB2r | R | GGAATTCGATATCAAGCTTGGC | RNAi inserts amplification |
| 1722 | F | ATCGCCAATGTGATGGggtattggatttgacggcgc | 8.4240 amplification for pQuadra (gene-specific part in lower case) |
| 1723 | R | TAGCCCATAGAGTTGGggtacgggctacacgtatgg | 8.4240 amplification for pQuadra (gene-specific part in lower case) |
| 116 | F | CTGCTGTGCCATCAGATTACTC | pQuadra sequencing primer 1 |
| 102 | F | TGCACGCGCCTTCGAGTT | pQuadra sequencing primer 2 |
| 82 | F | GAGTACTGAGTTTAACATGTTCTC | pQuadra sequencing primer 3 |
| 1777 | F | TGGCTCGAATGGTTGCTGGA | 8.4240 primers for qRT-PCR |
| 1778 | R | AAAGTGTTGTGATGCCCGCC | 8.4240 primers for qRT-PCR |
| 141 | F | CGGCGGCCGCATGTCAGGTAAACTTCGTCTTTACAAAG | L262P mutation confirmation |
| 6 | R | ATAGGATCCCTACTTGGTTACTGCCCCTTCCCAG | L262P mutation confirmation |
| 1358 | F | GAGCGTGTGACTTCCGAAGG | Tb927.11.10190 (TERT) primers for qRT-PCR |
| 1359 | R | AGGAACTGTCACGGAGTTTGC | Tb927.11.10190 (TERT) primers for qRT-PCR |
